# Stability through plasticity: Finding robust memories through representational drift

**DOI:** 10.1101/2024.12.19.629245

**Authors:** Maanasa Natrajan, James E. Fitzgerald

## Abstract

Memories are believed to be stored in synapses and retrieved through the reactivation of neural ensembles. Learning alters synaptic weights, which can interfere with previously stored memories that share the same synapses, creating a tradeoff between plasticity and stability. Interestingly, neural representations exhibit significant dynamics, even in stable environments, without apparent learning or forgetting—a phenomenon known as representational drift. Theoretical studies have suggested that multiple neural representations can correspond to a memory, with post-learning exploration of these representation solutions driving drift. However, it remains unclear whether representations explored through drift differ from those learned or offer unique advantages. Here we show that representational drift uncovers noise-robust representations that are otherwise difficult to learn. We first define the non-linear solution space manifold of synaptic weights for a fixed input-output mapping, which allows us to disentangle drift from learning and forgetting and simulate representational drift as diffusion within this manifold. Solutions explored by drift have many inactive and saturated neurons, making them robust to weight perturbations due to noise or continual learning. Such solutions are prevalent and entropically favored by drift, but their lack of gradients makes them difficult to learn and non-conducive to further learning. To overcome this, we introduce an allocation procedure that selectively shifts representations for new information into a learning-conducive regime. By combining allocation with drift, we resolve the tradeoff between learnability and robustness.

## Introduction

Cognition and behavior are thought to arise from neural activity patterns, leading many to equate brain functions with specific activity patterns, also called neural representations. For example, perhaps our sense of where we are amounts to which “place cells” fire in the hippocampus. Memory could in turn be the reactivation of neural activity patterns corresponding to past experiences, with Hebbian assemblies and memory engrams being prominent examples (1–3). However, recent data suggest that the mapping between neural representations and brain function is not one-to-one. In the hippocampal example, which neurons represent any given place changes over time (4–10). Such shifts in neural representations are termed “representational drift” (11, 12), because they appear to be devoid of discernible learning, forgetting, or behavioral changes. This intriguing phenomenon raises questions about the neural substrates of experience and its memory.

Relatively clear hypotheses exist for how neural representations could drift without disrupting cognition or behavior. In machine learning, researchers train artificial neural networks to generate desired input-output mappings, and neurons in intermediate layers can adopt any activity pattern that yields the correct output. The fixed inputs and outputs somewhat anchor the intermediate representations, but many possibilities remain. Drift can thus be modeled in neural networks as the random exploration of intermediate representations that produce the same performance (13–17). In biology, sensory and motor representations rigidly tie brain activity to the external world, and the brain’s cognitive functions may impose additional constraints. Yet it is unlikely that these anchors fully remove all the redundancies. Supporting this view, representational drift is seen across many brain regions, including the hippocampus (6), posterior parietal cortex (18), pre-limbic cortex (19), visual cortex (20), somatosensory cortex (21), and piriform cortex (22). However, drift seems to spare certain dimensions of neural activity (20, 23–28), pointing to low-dimensional substrates that might stably encode sensory, behavioral, and cognitive variables.

Drift may be a result of noisy learning systems. Many researchers have proposed that neural representations underlying memories are prone to degradation from factors like intrinsic noise and continual learning (29, 30). Error-corrective processes could compensate, but it’s likely that these will uncover a distinct representation than revert to the previous version, resulting in exploration of the solution space of the intact memory and representational drift (15, 17, 31).

Given that both learning and drift explore representations encoding the same memory content, a fundamental question arises. How might representations uncovered through drift differ from those initially learned? Moreover, could drift-induced representations offer advantages over the original learned solutions? Here we propose that drift benefits neural systems by helping them find noise-robust solutions. Our findings reveal that sparsely engaged representations, characterized by many inactive and saturated neurons, are more prevalent than densely engaged ones with fewer such neurons. Thus, random explorations in weight space favor these sparse representations for entropic reasons. Importantly, these common, sparse representations exhibit greater robustness to weight perturbations, which may occur due to noise or learning. However, these robust solutions are hard to learn, suggesting that drift may play a critical role in finding these solutions. Consequently, our results suggest that representational drift isn’t merely a byproduct of noise and continuous learning. Rather, it can actively help maintain stable memories in the face of noise and continual learning.

## Results

### Theoretical framework for modeling rep-resentational drift

We conceptualize representational drift as the free exploration of neural network states that result in the same behavior, by which we mean both cognitive processes and the control of the body. The neural mechanisms underlying behavior are incompletely understood, so here we model it as an abstract function, *Z*, of neural activity, *Y* (Fig. 1A). For example, *Z* might represent the visual perception of an apple, *Y* could denote the firing rates of neurons in the visual thalamus, and *Z*(*Y*) would be a complicated nonlinear function. Alternatively, if *Y* represents activity in high-level visual cortex, *Z*(*Y*) might be a simple linear readout. We assume that the neural activity *Y* is mechanistically determined by a synaptic weight matrix that combines recurrent inputs within *Y* and feed-forward drive from an input population, *X*. For example, *X* might be retinal ganglion ganglion cells if *Y* is visual thalamus. We ensure finite synaptic weights by imposing a weight norm bound for each postsynaptic neuron (SI Appendix SI Methods). We also assume that the nonlinear activation function relating input currents to firing rates, Φ, has finite activation and saturation thresholds. For instance, a neuron has a firing rate of zero when the input current is below its activation threshold, and a neuron’s refractory period imposes a limit on its firing rate. We refer to neurons with intermediate activity as “engaged,” since their activity changes with variations in input. In contrast, neurons that are either inactive or saturated do not alter their activity in response to input variations, and we refer to them as “disengaged.” In this work, we focus on a single drifting population by assuming that *X* is stable and *Z*(*Y*) is a fixed function unaffected by drift. Representational drift in *Y* thus results from changing synaptic weights, and we model it as diffusion in the solution space of synaptic weights that generate specified mappings between *X* and *Z* (Fig. 1B). We further assume that the number of input-output mappings, *P*, does not exceed the number of neurons providing input to *Y*. This allows us to locally construct a natural coordinate system for the solution space manifold (32, 33).

**Fig. 1.**
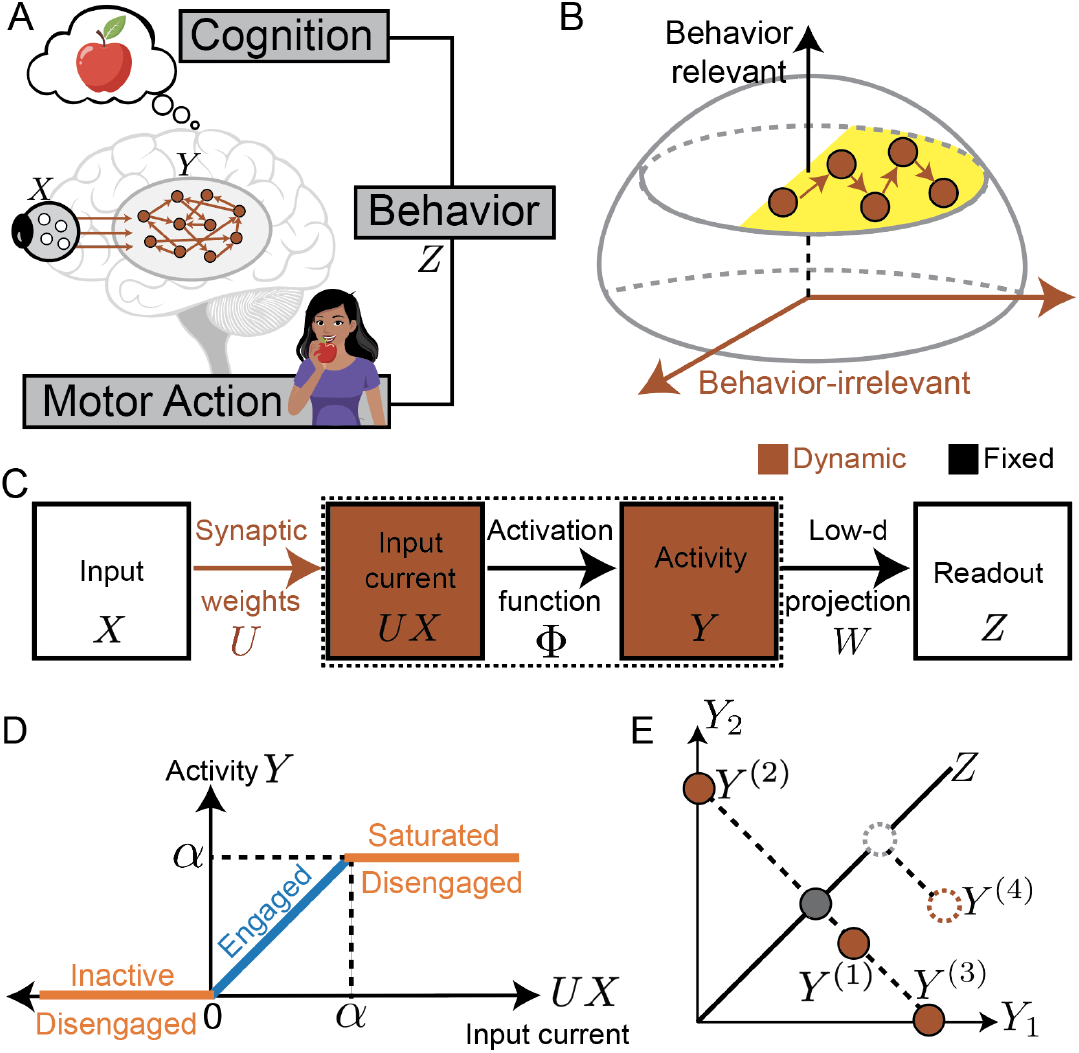
Theoretical framework for modeling representational drift: (A) Neural representation *Y* is generated from inputs *X*, and determines cognitive or motor readouts *Z*. (B) Illustration of a solution space (in yellow) of weights that generate the desired readout, and drift as a bounded diffusion restricted to the solution space. (C) Series of transformations that leads from input (*X*) to readout (*Z*), with components that can change during drift shown in brown and fixed components shown in black. Elements within the dashed border correspond to neurons participating in the analyzed representation (*Y*). (D) Depiction of the clipped-threshold-linear activation function that relates the input current (*UX*) to neural activity (*Y*). (E) Schematic showing behavioral readout (*Z*) as a low-dimensional projection of neural representation (*Y*), highlighting multiple representations may produce the same readout.

Most of our analyses will focus on a simple model variant that makes several additional assumptions (Fig. 1C). First, we ignore recurrent connections within *Y*. Thus, neural activity in the drifting population is a feed-forward transformation of *X* through synaptic weights *U*. Second, we model the firing rate nonlinearity as a clipped-threshold-linear function (Fig. 1D). This implies that neurons remain inactive when the input current is below 0, become saturated when the input current exceeds *α*, and linearly increase their firing rates in between. This function makes it straightforward to specify the solution space manifold, but the solution space can also be analyzed for other activation functions (SI Appendix A.1). Third, we assume that the behavioral readout *Z* is a 1-dimensional linear projection of *Y* (Fig. 1E). We can therefore specify *Z* through readout weights, *W*, but these weights are abstract and need not correspond to synaptic weights in the brain. This assumption is inspired by the empirical observation that behaviorally relevant information can often be linearly decoded from neural activity (34, 35). Nevertheless, nonlinear readouts may be required to understand representational drift in specific areas of interest (22–24, 36).

### Intuitive summary of framework and main results

Humans and other animals exhibit a diverse array of behaviors ranging from navigation to communication, which are generated in response to specific sensory cues and shaped by memories of past experiences. For example, to reach their lab a student might use a restaurant as a landmark to turn right and a library as a cue to turn left (37). Similarly, a monkey might emit a particular pitch vocalization to alert others of a leopard, while using different pitch sounds on seeing other predators (38). In each case, the behavior is a low-dimensional output represented and generated by collective neural activity. What ultimately matters is not the specific neural representation or synaptic weights, but generating the correct behavior. For example, the student needs to make the correct turns to reach their lab, and the monkey must produce the right vocalization to warn of specific predators. Although the system is constrained by these demands, if behavior *Z* is a low-dimensional projection of the neural representation *Y*, many different representations, and in turn many synaptic weights *U*, may produce the same output for given inputs (Fig. 2A) (11, 13). This implies a synaptic weight solution space that achieves the same behavior (Fig. 1B). In this paper, we will develop a framework for locally defining this solution space manifold and simulate drift as diffusion within it.

**Fig. 2.**
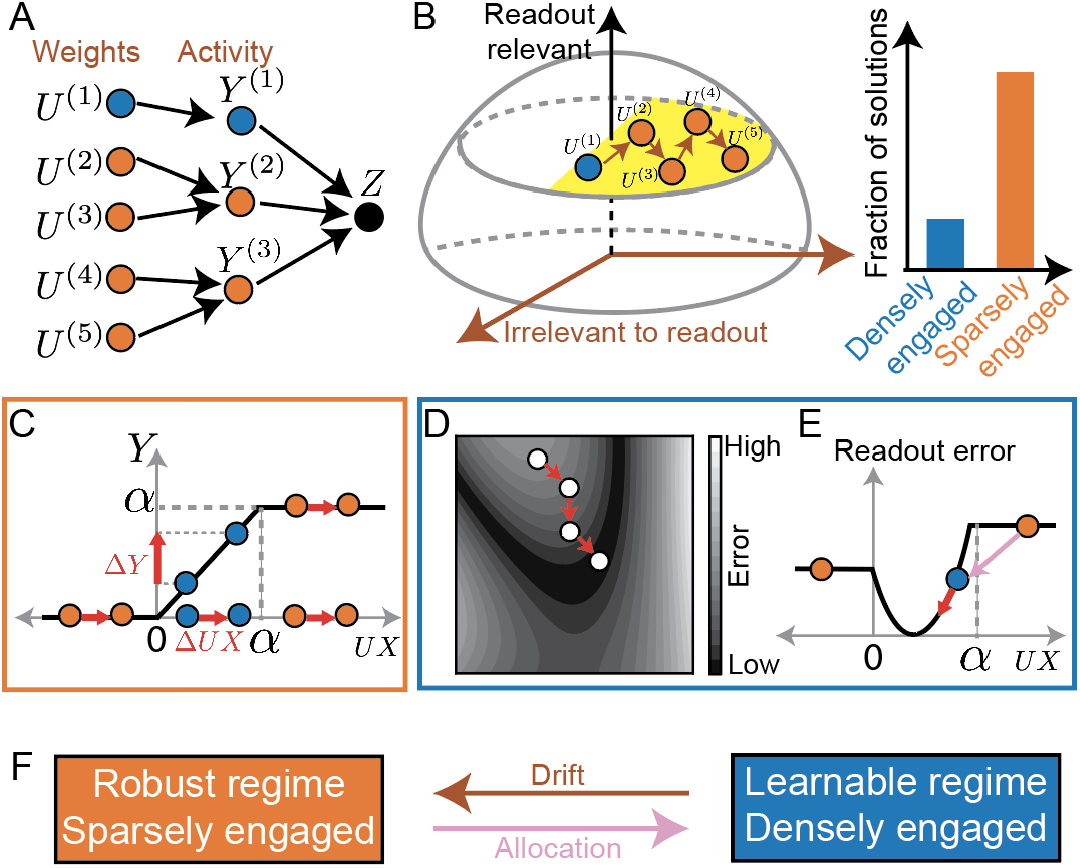
Drift as diffusion in the solution space improves robustness at the cost of learnability: An intuitive summary of main results. (A) Visualization of an example mapping between synaptic weight configurations (*U*), neural representations (*Y*), and readout (*Z*) with densely engaged solutions in blue and sparsely engaged solutions in orange. (B) Schematic indicating that sparsely engaged solutions may be more prevalent within the solution space. Unbiased exploration in the weight solution space (in yellow) can lead to sparsely engaged solutions due to their prevalence. (C) Illustration that disengaged neurons (inactive or saturated) enhance robustness to changes in input currents. (D) Diagram showing that learning requires movement in a direction opposite to the error gradient. (E) Illustration that engaged neurons provide gradient information, enabling learning. (F) Schematic showing how drift along with allocation may help overcome learnability-robustness tradeoff.

Diffusion in the solution space of synaptic weights implies that the system is equally likely to occupy any weight solution at a given time, so understanding representational drift requires a statistical description of the solution space. We first note that unbiased sampling of synaptic weight solutions can lead to biased sampling at the representation level. All non-positive input currents result in an inactive neuron, all input currents exceeding the saturation threshold produce a saturated neuron, but a single input current leads a neuron to be engaged with a specific activity level (Fig. 1C). We will show that this generates dimensions in which synaptic weights and input currents can change without changing the representation, and the number of such dimensions is set by the number of disengaged (inactive or saturated) neurons. If not for the weight norm bound, the system could diffuse infinitely far into these dimensions. When the weight norm bound is large, this leads sparsely engaged representations to be more common (Fig. 2A,B, SI Appendix B.1, Suppl. Fig. 3). Thus, we will see that representational drift can entropically favor sparsely engaged representations (Fig. 2B).

**Fig. 3.**
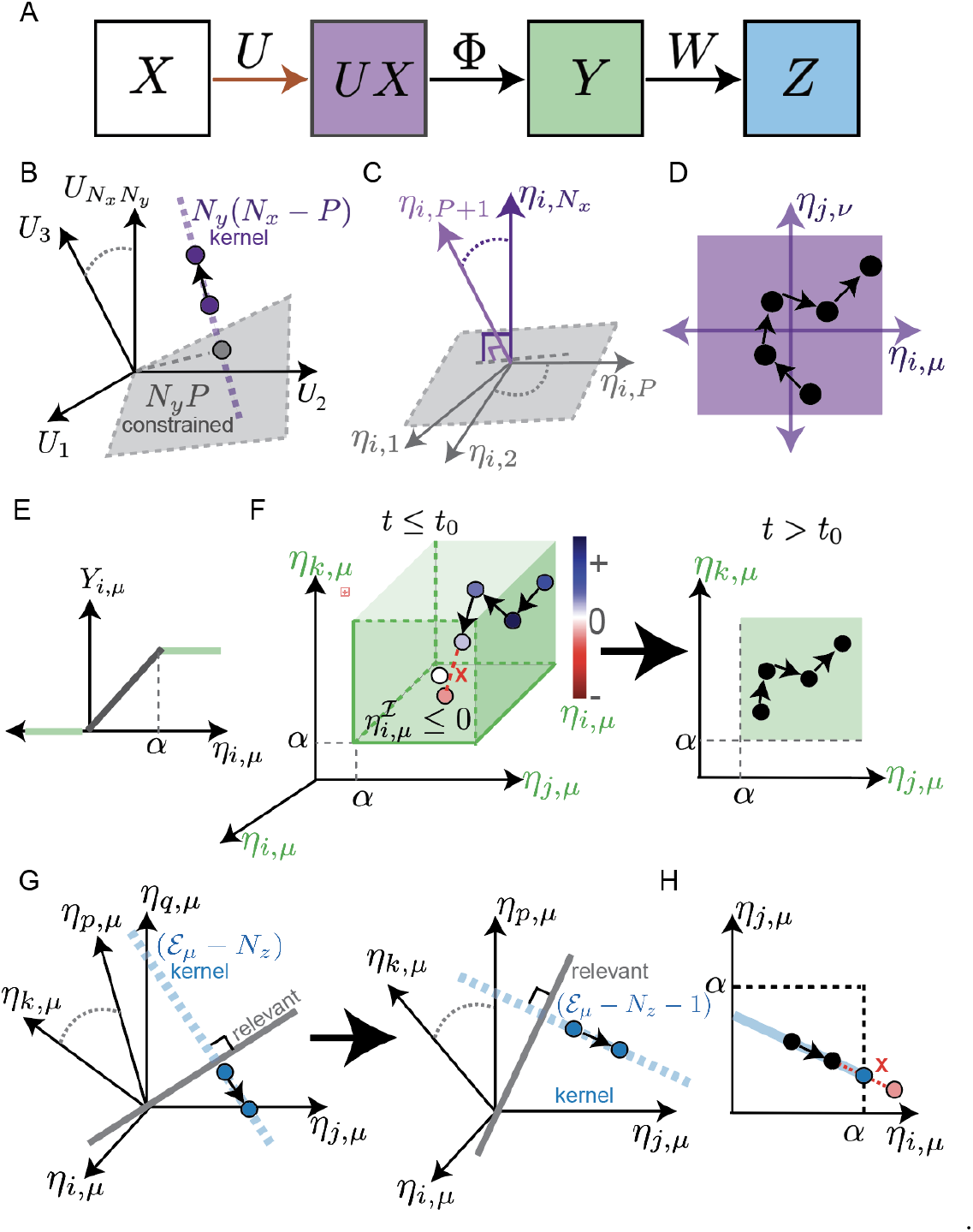
Characterization of the solution space and restricted diffusion. (A) Two linear-transformations, *U* and *W*, and one non-linear transformation, Φ, must together generate the behavioral readout *Z* from sensory input *X* via representation *Y*. Changing *U* (brown arrow) can result in fixed *UX* (purple, B-D), dynamic *UX* but fixed *Y* (green, E-F), or dynamic *Y* but fixed *Z* (blue, G-H). (B) Schematic depicting the *N*_*y*_ *P* - dimensional constrained subspace (gray), into which projections fully determine *UX*, and the *N*_*y*_ (*N*_*x*_ −*P*)-dimensional kernel of *X* (purple), in which unconstrained changes can occur. (C) Schematic of *η*-coordinates, which provide a convenient coordinate system on the synaptic weights by separating constrained (gray) and unconstrained (purple) dimensions for each neuron *i*. (D) Schematic of weight diffusion in unconstrained dimensions, where *µ > P, ν > P*, and *i, j* index arbitrary postsynaptic neurons. (E) The activation and saturation thresholds of the clipped-threshold-linear activation function provide flexibility by producing the same output for many inputs. This flexibility is confined to one side of the threshold, so we refer to its dimensions as semi-constrained. (F) Semiconstrained dimensions change over time. (Left) Diffusion in the semi-constrained subspace, where *µ* indexes the condition, neuron *i* is inactive, and neurons *j* and neuron *k* are saturated. At time *t*_0_, a random change would cause neuron *i* to cross the activation threshold and become engaged. *η*_*iµ*_ is set to 0, and neuron *i* is shifted to engaged set. (Right) After *t*_0_, the number of semi-constrained dimensions is reduced and the semi-constrained diffusion continues. (G) Representational drift dimensions change over time and involve multiple neurons. Visualization of the *E*^*µ*^-dimensional engaged subspace for condition *µ* ≤ *P*, with ℰ_*µ*_ = *{i, j, k, …, p, q}*. The engaged submatrix *W*_*µ*_ defines a *N*_*z*_ -dimensional relevant projection (gray), leaving *E*_*µ*_ − *N*_*z*_ flexible dimensions in its kernel (blue). (Left) Changes in these flexible dimensions cause representational drift, and neuron *q* eventually crosses the activity threshold and becomes inactive at time *t*_0_. (Right) In subsequent time points, the flexible dimensions are confined to the kernel of the new engaged submatrix, which has *E*_*µ,t*+_ − *N*_*z*_ = *E*_*µ,t*−_ − 1 − *N*_*z*_ dimensions, where *t*^−^ and *t*_+_ denote times shortly before and shortly after *t*_0_. (H) Schematic illustrating how random changes are rescaled to keep drift in the kernel of *W*^*µ*^ and prevent. threshold crossings, where *i, j* ∈ *ℰ*_*µ*_

Although sparsely and densely engaged solutions produce the same behaviors, sparsely engaged representations favored by drift are advantageous for other reasons. Returning to the example of a student navigating to their lab, consider two scenarios: first, the library used as a landmark becomes partially obscured by construction wrap, slightly altering the sensory input *X*. Alternatively, suppose the same neural circuit is used to learn the route to a coffee shop, leading to modifications in the weights *U*. In either case, the input current *UX* experiences a perturbation, but the student must turn correctly to reach their lab. Neural representations that produce the same behavioral output can respond quite differently to these perturbations. Disengaged neurons are robust to small changes in input currents because their activity remains constant unless the thresholds are crossed (Fig. 2C). Engaged neurons, on the other hand, are sensitive to changes in input currents, which can influence their activity and potentially alter the behavioral output. Thus, disengaged neurons contribute to robustness against noise and perturbations, making sparsely engaged representations more robust.

This robustness comes at a cost. While the exact plasticity rules that govern synaptic changes remain unclear, learning requires that synaptic modifications lead to performance improvements (Fig. 2D). For example, if a student typically goes straight upon seeing the library, but a change in synaptic weights causes them to turn right instead, they might reach their lab more quickly, leading to better performance. However, only engaged neurons are responsive to these synaptic weight changes and thereby able to affect neural activity and behavior (Fig. 2E). Engaged neurons, but not disengaged ones, provide feedback on performance changes. The very property that makes disengaged neurons good for robustness—resistance to perturbations—makes them less useful for learning.

This reveals a fundamental tradeoff between learnability and robustness. Sparsely engaged representations are good for maintaining stable memories; densely engaged representations are better for learning. We will see that representational drift naturally leads to sparsely engaged representations, facilitating a transition from learning-conducive states to more robust ones (Fig. 2F). However, to support new learning, we will also need a mechanism that shifts representations back to the learnable regime. We will thus introduce an allocation process that moves neurons into the engaged regime for new input conditions, creating a densely engaged representation conducive to learning, without disrupting previously stored information. By combining drift with allocation, we propose a dynamic solution to the learnability-robustness tradeoff in which the brain could maintain both adaptability for new learning and robustness for long-term memory retention.

### Characterization of the solution space and restricted diffusion

To model representational drift, we first develop a mathematical approach to simulate diffusion restricted to the solution space. We consider an *N*_*y*_-dimensional representation which has to generate an *N*_*z*_-dimensional readout from its *N*_*x*_-dimensional input under *P* different input conditions. The readout is modeled as a linear projection of the neural representation. The full input-output transformation of the system is thus

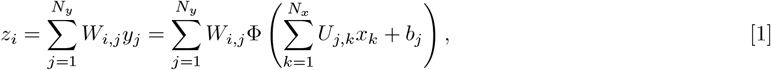

where *z* is an *N*_*z*_-vector defining the readout, *W* is a *N*_*z*_ *× N*_*y*_ matrix for the linear projection, *y* is an *N*_*y*_-vector for the representation, *U* is an *N*_*y*_ *× N*_*x*_ matrix of synaptic weights, *x* is an *N*_*x*_-vector of inputs, *b* is an *N*_*x*_-vector of biases, and Φ is the clipped-threshold-linear activation function with saturation threshold *α* relating input current to activity (Fig. 3A) (Eq. 2)

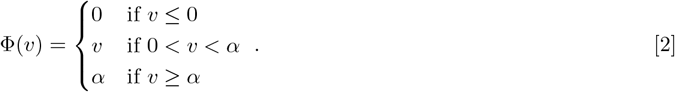

The constraints imposed by the *P* input-output mappings are

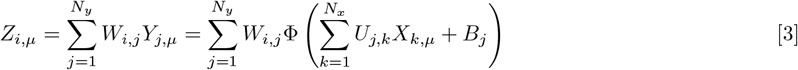

(Fig. 3A), where *µ* = 1, …, *P* indexes the conditions, *Z* is an *N*_*z*_ *× P* matrix of the required readouts, *Y* is an *N*_*y*_ *× P* matrix of representation across *P* conditions, *X* is an *N*_*x*_ *× P* matrix of inputs, and *B* is a *N*_*y*_ *× P* matrix formed by repeating bias vector *b* as columns *P* times. Note that Eq. 3 is simply *Z* = *W* Φ (*UX* + *B*) in matrix notation.

We model representational drift as diffusion in the solution space of weights *U* that satisfy Eq. 3 for *µ* = 1, *…, P*. Specifically, we characterize three kinds of flexibility: (1) changes in synaptic weights *U* that maintain the input currents *UX*; (2) changes in input currents that maintain the representation *Y* ; and (3) changes in the representation *Y* that maintain the readout *Z*. The first two kinds of flexibility entail synaptic weight changes concealed at the representation level, so we call these *covert exploration* dimensions. The third kind alters the representation, so we call these *drift* dimensions.

The first type of flexibility arises when the number of input neurons exceeds the number of conditions, *N*_*x*_ *> P*, because this implies a many-to-one mapping from weights to input currents. In particular, changes in *U* limited to the *N*_*y*_(*N*_*x*_ *−P*)-dimensional kernel of *X* don’t alter the input current *UX* (Fig. 3B). We call these unconstrained dimensions because there is complete freedom in these dimensions. Unconstrained dimensions can be separated out when we consider *U* in a coordinate system designed to manifest the activity-dependent constraints (Fig. 3C) (32, 33). We introduce a basis transformation matrix, *X*_*ext*_, where *X*_*ext*_ is an *N*_*x*_ *× N*_*x*_ full-rank matrix composed of *X* and *N*_*x*_ − *P* columns that span the kernel of *X*. The new coordinates are given by

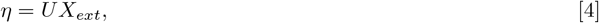

and we get *µ* = 1, *…, P* constraint equations,

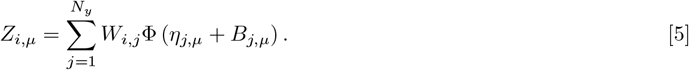

In these *η*-coordinates, the first *P* columns fully determine the input current *UX*. The last *N*_*x*_ − *P* columns are in the kernel of *X*, so they don’t affect the input current. Nevertheless, the synaptic weights depend on all columns of *X*_*ext*_ through

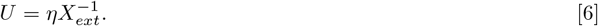

Note that this basis transformation acts independently on the incoming weights to each postsynaptic neuron. In order to make changes in the unconstrained dimensions, we make changes in *η* for all input conditions *µ > P* according to 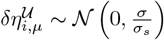 (Fig. 3D), where *σ* sets the scale of the weight fluctuations, and 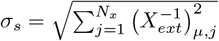 is chosen to generate equal variance weight fluctuations.

The second kind of flexibility arises due to the clipped-threshold-linear activation function, as changes in input currents don’t affect activity if the input currents remains below the activity threshold or above the saturation threshold (Fig. 3E). Since these conditions lead to inequality constraints that restrict the solution space to part of a dimension, we refer to these as semi-constrained dimensions (32, 33). Importantly, which dimensions are semi-constrained can vary over time, as semi-constrained dimensions obtain new constraints when inactive or saturated neurons become engaged. Thus, diffusion in semi-constrained dimensions interacts with drift dimensions.

To coordinate changes in semi-constrained dimensions with representational drift, we divide neurons into sets of engaged (ℰ ^*µ*^), inactive (ℐ^*µ*^), and saturated (𝒮^*µ*^) neurons for each input condition *µ* ≤*P*. These sets depend on time, but we will use elaborated notations like ℰ^*µ,t*^ only when it is important to emphasize this time dependence. The distinction between these sets is based on where the input current lies in relation to the thresholds, with neurons below the activity threshold (0) being inactive, neurons above the saturation threshold (*α*) being saturated, and neurons in between thresholds being engaged.

Neurons transition between these sets when their input current exactly corresponds to a threshold. For instance, for each neuron in the inactive set, we change its *η*-coordinates such that 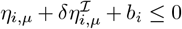, where *i* ∈ *ℐ*^*µ*^. However, when a random change, 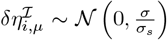, would have caused the input current for neuron *i* to cross its activity threshold at 0, we set the new *η*_*i,µ*_ to −*b*_*i*_ and shift neuron *i* from the inactive to engaged set (as in Fig. 3F). Similarly for neurons in the saturated set, we ensure that 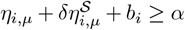, where *i* ^*µ*^, and when the input current would have crossed the saturation threshold at *α*, we set the new *η*_*i,µ*_ to *α − b*_*i*_ and shift *i* from saturated to engaged set. These transitions between semi-constrained and engaged dimensions play a critical role in changing the sparsity of solutions explored by drift.

Diffusion in the representational drift dimensions typically requires concerted changes between multiple neurons in the population. By definition, this entails changes in representation that are orthogonal to the readouts. This implies that drift must be in the kernel of *W*. However, since small weight changes do not affect the activity of inactive and saturated neurons, drift can only involve changes to the activity of neurons in the engaged set. Therefore, representational drift must locally occur in the kernel of the *engaged submatrix, W* ^*µ*^, which only contains the columns of *W* that correspond to engaged neurons (Fig. 3G, left). In particular, since there are *N*_*z*_ readout dimensions and *E*^*µ*^ = |ℰ^*µ*^| engaged neurons for input *µ*, drift changes are within the (*E*^*µ*^ − *N*_*z*_)-dimensional kernel of *W* ^*µ*^. As the engaged set for each input condition changes during drift, the engaged submatrix, the dimensionality of drift, and the dimensions of drift change with time. We will use the more explicit notations of *E*^*µ,t*^ when useful.

We sample drift changes as 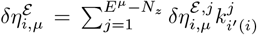, where 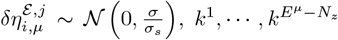 represent orthonormal basis vectors of ker *W* ^*µ*^, and *i*^*′*^ : ℰ^*µ*^ →{1,…, *E*^*µ*^} is simply a function that converts indices between *N*_*y*_-dimensional vectors and *E*^*µ*^-dimensional vectors by replacing *i* with its position in ℰ ^*µ*^. For example, if ℰ^*µ*^ = {1, 10, 12}, then *i*^*′*^(1) = 1, *i*^*′*^(10) = 2, and *i*^*′*^(12) = 3. Due to the clipped-threshold-linear activation function, activity is bound between 0 and *α*, which imposes inequality constraints on representational drift akin to those for the changes in semi-constrained dimensions, 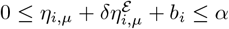, where *i ℰ*^*µ*^. However, if a neuron would have crossed one of the thresholds, we cannot simply set its *η*-coordinate to the threshold, since that change would no longer be in the kernel of *W* ^*µ*^. Instead, we rescale the magnitude of the change so that the neuron sits exactly at threshold (Fig. 3H) (see Materials and Methods). When more than one neuron would have crossed threshold, the rescaled change is the maximal that ensures that no neurons cross threshold. In either case, the neuron that reaches threshold is shifted from the engaged set to the inactive or saturated set depending on which threshold was reached. For instance, in Fig. 3G one neuron reaches the 0 threshold and becomes inactive. At the next time point, there is one fewer engaged neuron, and drift is limited to a lower-dimensional kernel.

By combining *δη*^*𝒰*^, *δη*^*ℐ*^, *δη*^*𝒮*^, and *δη*^*ℰ*^, we get *δη* as

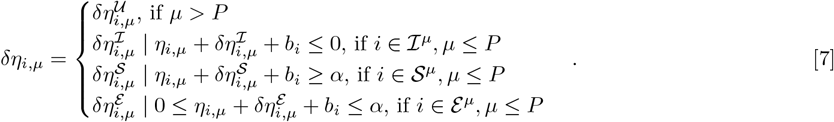

We compute the new *η*-coordinates as *η* + *δη* and transform back to the physical coordinates using Eq. 6. This method implements diffusive changes in *η*-coordinates, but the resulting *δU* is approximately normal with mean 0 and standard deviation *σ* in physical coordinates (SI Suppl. Fig. 1).

### Allocation and learning without interference

Beyond defining the solution space, *η*-coordinates also enable targeted allocation and learning without interference. To understand these processes, consider a toy example with a two-neuron representation (*y*_1,*µ*_, *y*_2,*µ*_) that receives input from (*x*_1,*µ*_, *x*_2,*µ*_) via synaptic weights *U* and needs to generate a 1-dimensional readouts *z*_*µ*_ for each of the 2 mappings (*µ* = 1, 2) to be stored (Fig. 4A). We use readout weights *w* = (1, 1), no bias (*b* = (0, 0)), and a clipped-threshold-linear activation function Φ (Eq. 2) with activity and saturation thresholds at 0 and 5 respectively. Instead of directly visualizing the 4-dimensional weight matrix *U*, we will use the basis transformed *η*-coordinates defined as

**Fig. 4.**
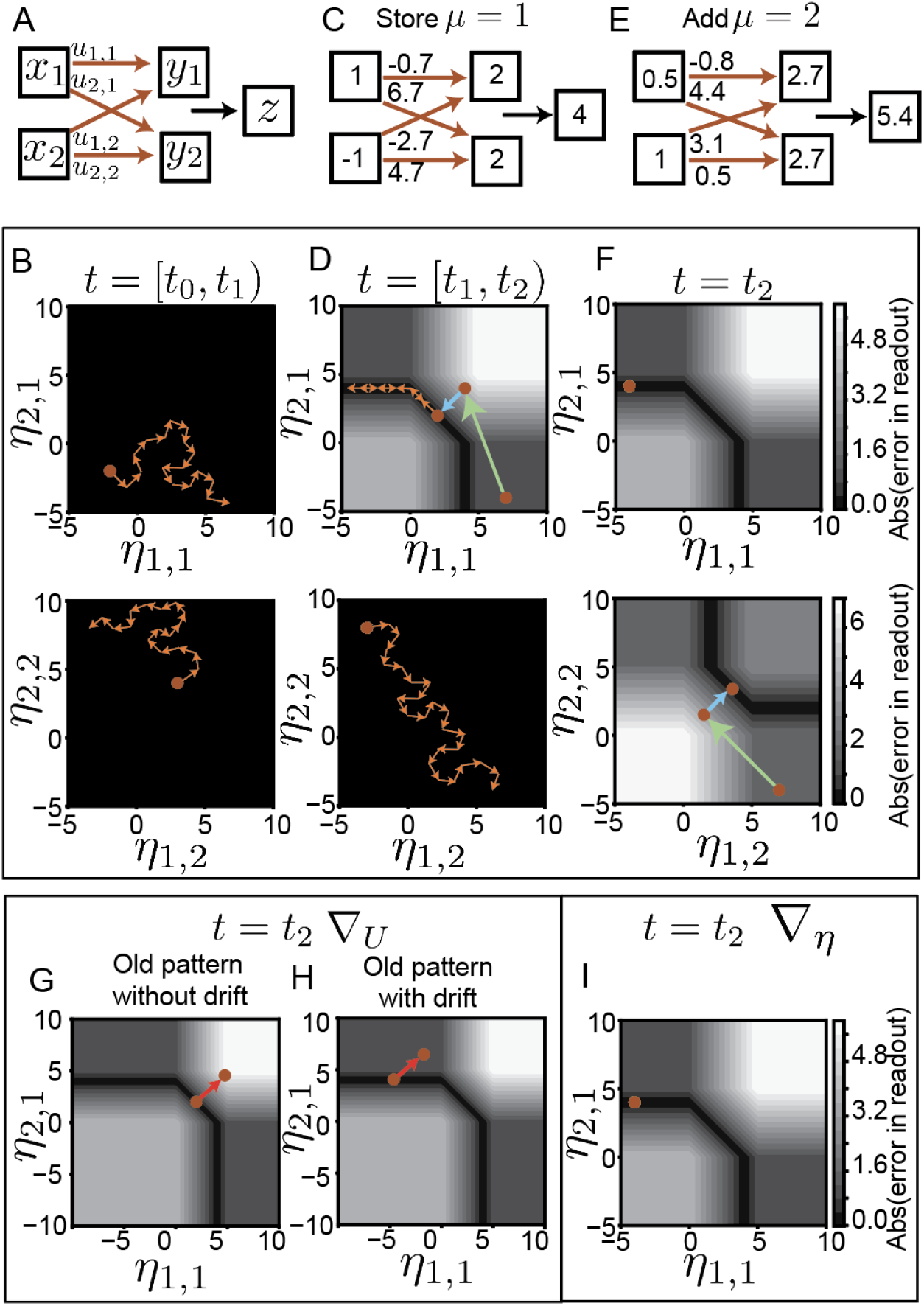
A toy example illustrating combining allocation, learning, and representational drift. (A) Network architecture for a toy example, with two inputs, a representation-layer with two neurons, and a single readout *z* with fixed readout weights *w* = (1, 1). (B,D,F) The 4-dimensional synaptic weights are visualized using a basis transformed version *η* based on input patterns that will be added to the network. Input currents corresponding to the first pattern *η*_*µ*=1_ are shown at the top and that corresponding to the second pattern *η*_*µ*=2_ at the bottom. (B) From time [*t*_0_, *t*_1_), no memories are added to the network, and *η* (brown point) diffuses freely through drift (orange arrows). (C, D) At *t* = *t*_1_ a new input-readout mapping (*µ* = 1) from (1, − 1) to (4) is to be stored. The contour plots indicate the error surface in *η*_*µ*=1_ (top). Green arrow denotes the allocation of *η*_*µ*=1_ to a dense representation (4, 4) while the blue arrow shows gradient descent performed in *η*_*µ*=1_ to reach the solution space at (2, 2). *η*_*µ*=2_ remains unchanged during allocation and learning (bottom). Between *t*_1_ and *t*_2_ representational drift continues, with exploration constrained to the solution space in *η*_*µ*=1_ and free in *η*_*µ*=2_ coordinates (orange arrows). (E, F) At *t* = *t*_2_, a new input-readout mapping (*µ* = 2) from (0.5, 1) to (5.4) is to be added, generating an additional error surface (bottom) on *η*_*µ*=2_. Allocation (green arrow) followed by gradient descent (blue arrow) are restricted to *η*_*µ*=2_. (G,H) Gradient descent in *U* to learn *µ* = 2, error contours with respect to *µ* = 1. (G) Brown point indicates the input currents due to learning of *µ* = 1. Red arrow denotes the change in *η*_*µ*=1_, due to the addition of new mapping *µ* = 2, resulting in large error for *µ* = 1 mapping. (H) Brown point indicates the input currents corresponding to *µ* = 1 after learning followed by drift (sparser solution). Red arrow indicates the change in *η*_*µ*=1_ due to addition of new mapping, resulting in a comparatively lower readout error for *µ* = 1 due to the enhanced robustness of the post-drift sparsely engaged solution. (I) Gradient descent in *η*-coordinates to learn *µ* = 2, error contours with respect to *µ* = 1 showing no change in input currents for the previously learned memory.

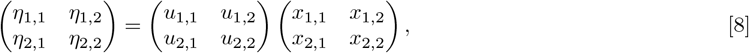

where *η*_*µ*=1_ = (*η*_1,1_, *η*_2,1_) and *η*_*µ*=2_ = (*η*_1,2_, *η*_2,2_) are weight combinations relevant for each of the two input conditions, *x*_*µ*=1_ = (*x*_1,1_, *x*_2,1_), and *x*_*µ*=2_ = (*x*_1,2_, *x*_2,2_), respectively.

*η*-coordinates enable targeted allocation facilitating learning. Between time *t*_0_ and *t*_1_, with no stored memories, *η* freely explores the synaptic parameter space via drift (Fig. 4B). At *t* = *t*_1_, when the system needs to learn a mapping from *x*_*µ*=1_ to *z*_*µ*=1_ (Fig. 4C), an error surface is introduced. The *η*-coordinates isolate the input-relevant weight combinations, so adding the first memory imposes an error surface on *η*_*µ*=1_ leaving *η*_*µ*=2_ with no constrains (Fig. 4D). Due to drift, the system is initially likely to be in a local optimum with both neurons disengaged for *µ* = 1. In the local optimum, errors don’t change locally, so without gradients, it’s unclear how to adjust weights for learning. To resolve this, we allocate the relevant weight combination, *η*_*µ*=1_, to random values centered in the engaged regime, leaving the weight combinations relevant to other inputs (*η*_*µ*=2_) unaffected (Fig. 4D). This moves the system into a densely engaged, gradient-rich learnable regime.

Drift subsequently enhances the robustness of learned memories. For *µ* = 1, starting from a densely engaged allocation, learning follows gradients to a nearby solution that is also densely engaged and thus non-robust. Before adding memory *µ* = 2, we allow the system to drift (Fig. 4D). During drift, it freely explores *η*_*µ*=2_ and diffuses within the *η*_*µ*=1_ solution space. Through drift, it finds more prevalent, sparsely engaged, robust solutions.

At *t* = *t*_2_, we aim to add a new mapping *µ* = 2 (Fig. 4E). This creates an error surface in *η*_*µ*=2_, while the error surface in *η*_*µ*=1_ remains unaffected (Fig. 4F) because only *η*_*µ*=2_ include weight combinations relevant to *µ* = 2. Again, we find that the configuration after drift generates representations with both disengaged neurons for *µ* = 2, trapping it in a local optimum. As before, we can enable learning by allocating *η*_*µ*=2_ to a densely engaged learnable regime.

Learning is often modeled as gradient descent on the error function. Let (*J*_2_)^2^ represent the squared readout error for *µ* = 2, (*J*_*2*_)^2^ = (Σ_*i*_ *w* Φ (Σ_*j*_*u*_*i,j*_ *x*_*j*,2_ − *z*_2_)^2^, where *z*_2_ is the required readout for *µ* = 2. Then the gradient of (*J*_2_)^2^ with respect to the weights indicates the direction of steepest increase in loss for *µ* = 2. To minimize the error, we update the weights in the direction opposite to the gradient, using

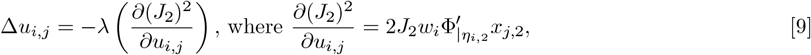

Where 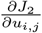 is the gradient, and *λ* is the learning rate.

Drift helps reduce interference due to new learning. Updating weights based solely on (*J*_2_), the error for *µ* = 2, is likely to disrupt the weight combinations relevant to the previously stored memory (*µ* = 1). This interference exemplifies the continual learning problem. The severity of disruption depends on the robustness of stored memories. Densely engaged representations are less robust, and thus more disrupted (Fig. 4G), while drifted memories, being sparsely engaged, remain robust to these weight updates (Fig. 4H).

Interference due to new learning can be completely eliminated using *η*-coordinates. In the *η* space, weight combinations relevant to each input condition are naturally segregated, so a new memory *µ* imposes its error surface only on its corresponding *η*_*µ*_. Therefore, learning implemented as gradient descent on the synaptic weights in *η*-coordinates only modifies the weight combinations relevant to the new memory,

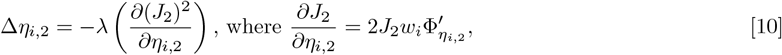

while preserving those associated with previously stored memories Δ*η*_*i*,1_ = 0 (Fig. 4I). When the number of input neurons exceeds the number of memories (*N*_*x*_ *> P*), *η*-space learning circumvents the continual learning problem, enabling sequential addition of memories without the need for batch learning. However, the *η*-coordinates of the weights are defined using a basis transformation matrix *X*_*ext*_ that incorporates all input conditions, leaving the biological implementation unclear (see Discussion).

### Unbiased exploration of weight solution space favors sparsely engaged, robust solutions

We simulated representational drift and studied the properties of the resulting representations. As anticipated, the representations changed (Fig. 5A,B) while the low-dimensional readouts remained stable (Fig. 5C). During the initial drift, the number of engaged neurons decreased, leading to sparsely engaged representations (Fig. 5D). Throughout later drift, the fraction of engaged neurons remained stably low, although the specific neurons in the engaged set continued to change (Fig. 5E). Under weight perturbations, sparsely engaged solutions explored by drift yielded smaller readout errors than the learned solutions (Fig. 5F). Thus, uniform sampling of the synaptic weight solution space resulted in a biased sampling of sparsely engaged, noise-robust solutions.

**Fig. 5.**
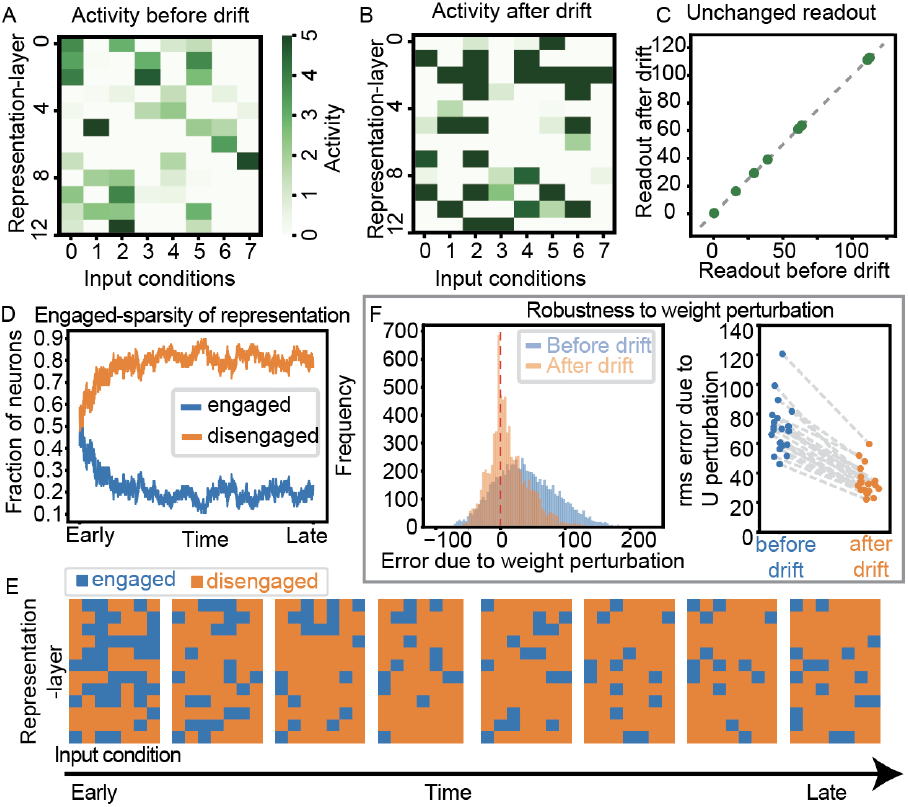
Representational drift in higher dimensions leads to sparsely engaged representations that are robust to weight perturbations. (A) Neural activity of 12 neurons at 8 different input conditions found using gradient descent learning. (B) Neural activity after simulating drift as a diffusion in the solution space. (C) Readout or 1-dimensional projection of the representation before and after drift, showing that it remains the same. (D) Fraction of engaged and disengaged (inactive and saturated) neurons over time during drift showing an initial drop in the fraction of engaged neurons followed by stable maintenance. (E) Engaged and disengaged neurons for each of the input conditions over time during drift. (F) Histogram of signed errors in readout due to *U* perturbations ∼ *𝒩* (0, 1). Root-measure squared error due to weight pertrubations before drift (blue) and after drift (orange)

### Representational drift increases robustness at the expense of learnability

Representational drift caused an increase in weight magnitudes. To ensure finite weights during drift, we imposed a weight norm bound *U*_*max*_ for each post-synaptic neuron *j*, 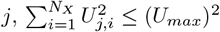 (Fig. 6A). In high-dimensional spheres, most of the volume is concentrated near the surface, meaning there are more weights with norms close to the imposed bound than smaller norms. Thus, drift entropically favored solutions with weight norms close to *U*_*max*_. Further, *U*_*max*_ greatly exceeded the weight norms of the initially learned solution, so during drift, weight norms increased to reach *U*_*max*_ and fluctuated close to it (Fig. 6B). Consequently, the synaptic weights post-drift were substantially larger than those prior to drift (Fig. 6C).

**Fig. 6.**
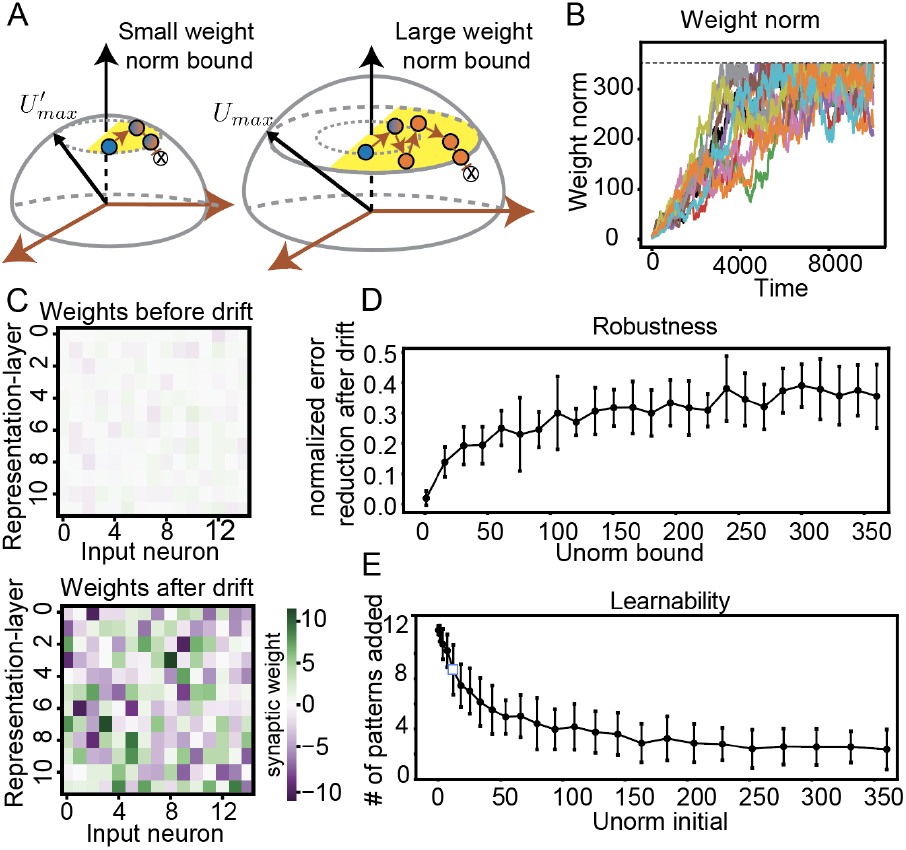
Representational drift increases robustness at the expense of learnability. (A) Schematic illustration of a small weight norm bound (based on initial weight norm) and a large weight norm bound, highlighting that densely engaged non-robust solutions (blue) are more prevalent for small weight norms, while sparsely engaged, robust solutions (orange) become more common for large weight norms. (B) Weight norms onto each post-synaptic neuron (depicted in different colors) over time during drift, showing an increase and plateau in weight norms close to the enforced weight norm bound, indicated by the black dotted line. (C) Small magnitude synaptic weights learned by gradient descent and large magnitude synaptic weights after representational drift. (D) Robustness: Difference between error in readout caused by random weight perturbations before and after drift, scaled by the error due to perturbation before drift, serving as a metric of the robustness gained due to drift. Different weight norm bounds during drift are considered. (E) Learnability, i.e., the number of input-readout mappings successfully stored in a network with varying initial weight norms. The graph demonstrates a decrease in learnability with an increase in initial weight norms

Large weight solutions explored by drift enhanced robustness. Higher weight magnitudes generate larger input currents, causing many to fall below the activity threshold or exceed the saturation threshold, leading to more disengaged neurons (SI Appendix B.2). The greater the number of disengaged neurons, the greater the robustness to weight perturbations (SI Appendix B.3). As a result, the robustness benefit from drift increases with the weight norm bound (Fig. 6D).

Large weight solutions explored by drift hindered future learning. After storing a few memories and allowing drift, the system must continue learning new input-readout mappings. However, large initial weights generate large input currents for new inputs, leading to sparsely engaged representations. This causes the gradients to vanish, preventing the system from learning new information. Consequently, the ability to learn new mappings decreased as initial weight magnitudes increased (Fig. 6E). This creates a tradeoff between retaining learned information through robust solutions and the capacity to learn new information, with larger weights favoring robustness but limiting learnability (Fig. 6D,E).

### Combining drift with allocation overcomes the learnability-robustness tradeoff and enables continual learning

Targeted allocation improves learnability despite large weight magnitudes. We initialized the network with large synaptic weights (*U* ∼ *𝒩* (0, 15)) and incrementally added new memories *µ* = 1, *…, P*. For each memory *µ*, we first allocated the relevant weight combinations *η*_*µ*_ ∼ *𝒩* (2.5, 0.1), near the center of the engaged regime (0,5). Next, we performed gradient descent learning in *η*_*µ*_, modifying only weight combinations relevant to the *µ*-input. Before learning the next pattern, we allowed the system to drift while preserving readouts for all learned patterns 1, *…, µ*. Using this approach, we successfully incorporated *N*_*x*_ memories, even slightly surpassing the learnability achieved with small weight initializations (Fig. 7A). While small weight initializations avoid saturation, they are equally likely to generate both inactive and engaged neurons. In contrast, the allocation method centers input currents within the engaged regime, ensuring densely engaged initializations (Fig. 7B), enhancing gradient availability, and thus learnability.

**Fig. 7.**
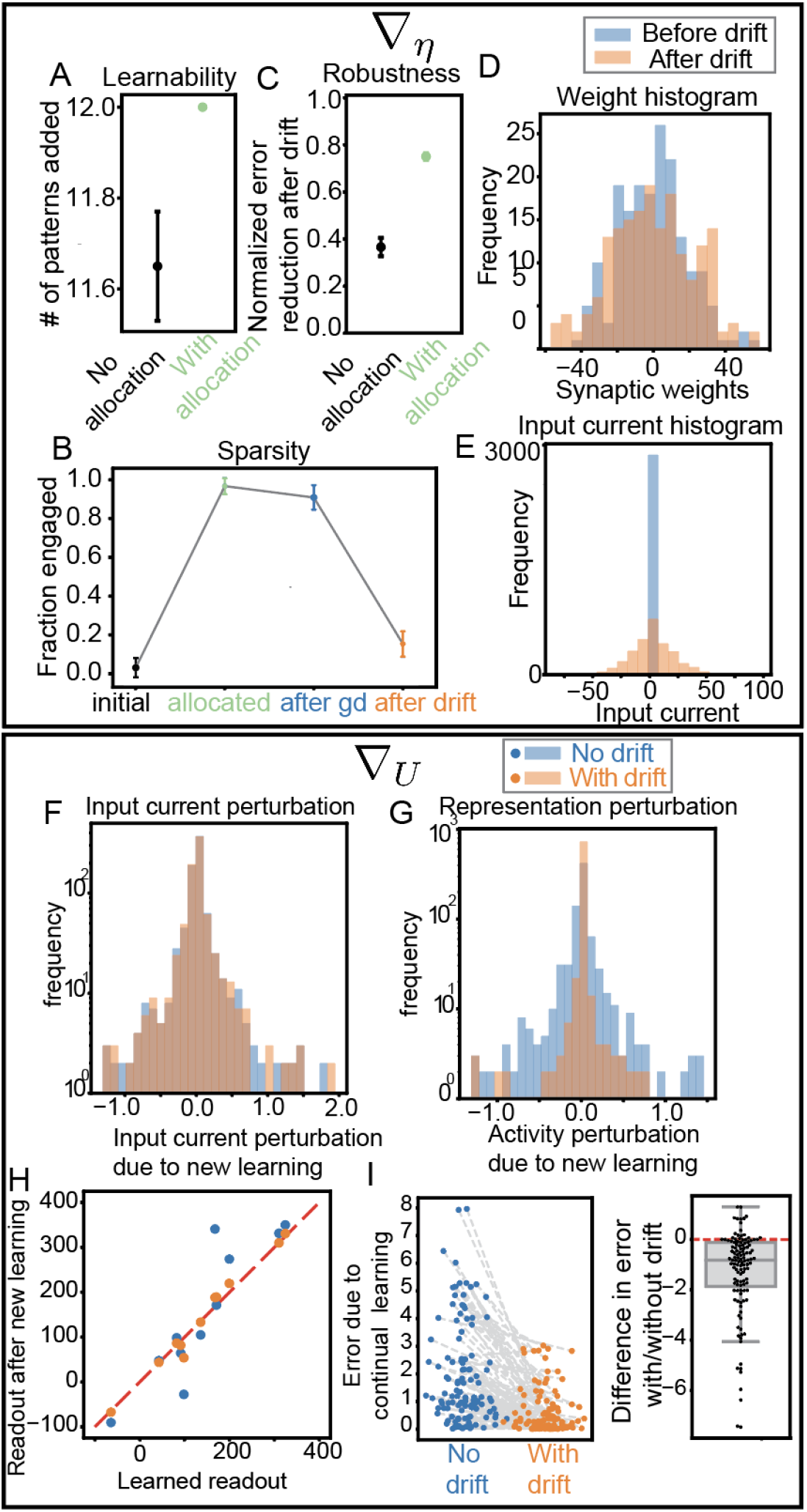
Allocation and representational drift effectively address the learnability-maintainability tradeoff and the continual learning problem. (A) The mean and standard error of the number of input-readout mappings that were successfully added to the network, with and without allocation. Without allocation shows the best learnability case from (Fig. 6E) with weight norm = 1. (B) Fraction of engaged neurons at different stages: initial (before allocation), after allocation, after gradient descent has converged, and after representational drift. (C) Difference between error in readout caused by random weight perturbations before and after drift, scaled by the error due to perturbation before drift, serving as a metric of the robustness gained due to drift. Without allocation (best robustness increase case from (Fig. 6D), with weight norm = 350) and with allocation. (D) Histogram of weights *U* before and after drift showing weights remain comparable (E) Input currents before and after drift showing drift causes input currents to spread out.(F,G,H,I) Continual learning simulations performing gradient descent in *U*, without drift (blue) and with drift (orange) in between to evaluate the effect of learning new pattern *ν* on previously stored patterns *µ*. (F) Histogram of perturbations to input currents *η*_*µ*_ due to new learning *ν*, showing same effect regardless of drift. (G) Histogram of perturbation to neural representation *Y*_*µ*_ indicating larger changes without drift compared to with drift(G). (H) For each pattern *µ*, readout upon learning *µ* vs readout after learning new pattern *ν*, with *µ < ν* ≤ *N*_*x*_. (I) Error in readout for pattern *µ* after learning of newer pattern *ν* and the difference between readout errors with drift and without drift, highlighting reduced errors with drift compared to without drift.

Drift can sparsify and enhance robustness without increasing weight magnitudes. Starting from densely engaged initializations, gradient descent caused some neurons to disengage, but the learned solutions remained densely engaged (Fig. 7B), and thus non-robust to weight perturbations. Drift reduced errors from perturbations, achieving greater robustness (Fig. 7C) while maintaining similar weight distribution (Fig. 7D), as the imposed weight norm bound was comparable to the learned weight norm. Increased robustness resulted from sparsification. The allocation procedure made the representations more densely engaged than typical for the set weight norm bound. During drift, the system explored more typical solutions with larger input currents (Fig. 7E), resulting in sparsely engaged (Fig. 7B), robust solutions (Fig. 7C).

Drift improves robustness to continual learning. We incrementally added new memories *µ* = 1, *…, P* by allocating *η*_*µ*_ and performing gradient descent on the synaptic weights *U* for the new memory *µ*. In one case the system drifted between memory additions, while in another memories were added without drift. In both cases, new learning caused the same input current perturbations to previously stored memories (Fig. 7F). However, because drift found sparsely engaged representations, the same input current perturbations had less impact on the neural representations (Fig. 7G) and, consequently, the readouts (Fig. 7H), resulting in lower readout errors (Fig. 7I). Thus, by increasing robustness to weight perturbations, drift improves memory maintenance during new learning.

In summary, learning and memory maintenance benefit from different representational regimes, and allocation and drift facilitate transitions between them. Densely engaged representations facilitate learning by generating gradients critical for learning, while sparsely engaged representations enhance memory stability by minimizing the impact of weight changes on readouts. *η* allocation boosts learnability by ensuring sufficient local gradients for new patterns to be learned, while drift safeguards existing memories by favoring robust, sparsely engaged solutions. Together, these processes support the learning of new information while stably preserving existing knowledge.

## Discussion

### The purpose of representational drift

Representational drift has been proposed to improve learning and memory in many ways. For instance, drift may help revise memories in light of new experiences (12, 39), or provide a baseline level of synaptic flux that helps overcome the exploration-exploitation dilemma in dynamic environments (15). Drift can prevent the system from getting stuck in local minima (11, 40) or improve generalization through drop-out regularization (26, 41). Other interesting proposals include probabilistic inference (16, 31), time stamping of memories (7), systems consolidation (39, 42), and memory decay to prevent catastrophic forgetting (12). Our theory provides interesting new insights on role of drift in overcoming the plasticity-stability dilemma. Many previously proposed advantages arise from enhanced exploration of different memories in dynamic environments that demand memories to change in order to align with the current context. Our work shows that representational drift can also provide benefits when memories are stably maintained, as it can mitigate interference caused by noise and continual learning. This provides an intriguing perspective on how representational drift relates to continual learning. Many have suggested that representational drift is a byproduct of error-corrective mechanisms that bring the system back to the solution space following perturbations caused by noise and continual learning (11, 15, 41).

Since we show that representational drift enhances robustness to noise and continual learning, this suggests that representational drift may fortify the system against the very processes that give rise to it. Our proposal that drift benefits continual learning through enhanced stability likewise offers a novel perspective on the stability-plasticity dilemma. Traditionally, stability is considered crucial for retaining stored memories and plasticity essential for learning new information. Thus, the stability-plasticity dilemma arises from the competing demands to balance memory maintenance and new learning (43, 44). In contrast, our results challenge the notion of a direct competition between plasticity and stability. Instead, plasticity-induced drift may actually bolster stability.

### Disentangling drift from learning

Our work demonstrates the power of mathematical modeling in disentangling representational drift from learning—two highly interwined processes. For instance, if representational drift results from ongoing learning (11), then the same biological mechanisms underlie both, making it experimentally challenging to isolate them. Nevertheless, here we mathematically isolated drift by modeling it as weight space diffusion in readout-irrelevant dimensions, analytically identified by extending prior theoretical work on the geometry of neural network solution spaces (32, 33). With perfect task performance and drift-induced changes orthogonal to the required readouts, we could isolate the benefits stemming solely from random exploration of the solution space. Moreover, this abstraction allowed our model to remain agnostic to what drives drift, whether that be ongoing learning, stochastic synaptic changes (15, 45, 46), spiking noise (47), noisy gradients (48), or fluctuating neuronal excitability (49).

This separation revealed that several phenomena typically associated with learning can also result from representational drift of stably maintained memories (SI Appendix C). For instance, stored memories are often correlated initially, but their representations decorrelate over time (50–52). This decorrelation reduces interference and improves separability (52–55), and it has often been attributed to learning. However, our work suggests that drift can also help orthogonalize memories. In high dimensional spaces, two randomly chosen vectors are close to orthogonal (56). Therefore, even if representations for two stimuli are initially correlated-due to encoding in temporal proximity (57, 58) or input similarity- representational drift is likely to decorrelate them over time (SI Appendix C.3, Suppl. Fig. 8). Similarly, phenomena such as representational sparsification (SI Appendix C.2, Suppl. Fig. 7) (41) and changes in representational turnover rate (SI Appendix C.1, Suppl. Fig. 6)(22), often associated with learning, may emerge from drift alone due to the geometry of solution spaces.

### Biologically plausible mechanisms for allocation

This paper considered a theoretically convenient yet highly abstract model of allocation, and it is critical to relate the three core elements of the model to candidate underlying mechanisms.

First, our model performs allocation prior to discrete learning events, but the triggers for allocation in biological systems might be more complex. Our results suggest that allocation may be necessary to encode new input-output mappings, so novelty detection could trigger allocation (59) especially when there are new sensory-behavior associations (60). More generally, allocation could be driven by whatever mechanisms the brain uses to prioritize information for memory encoding (61, 62). Since we proposed allocation to prevent learning from getting stuck in local optima, it could be beneficial if allocation is triggered by failed learning events. However, it’s unclear whether the brain detects such failures.

Second, our model implements allocation by resetting synaptic weights to values that provide a gradient signal, but this is only one of several mechanisms that biology could use. Directly instantiating our memory allocation model in biology would require input-dependent mechanisms that coordinate synaptic weight changes across many synapses. This is not inconceivable, as the required shifts can be subtle, and homeostatic plasticity can rescale many synapses onto a postsynaptic cell (63) and selectively modify activated synapses (64–66). Nevertheless, a more attractive alternative might involve transiently altering neuronal excitability, a mechanism widely associated with allocation in biology (67, 68). In our modeling framework, changes in excitability could correspond to changes in activity and saturation thresholds, thereby increasing neuronal engagement and providing gradients for learning. These changes could be made transient by homeostatically restoring the original thresholds in concert with compensatory weight changes to preserve relevant readouts.

Third, our model used the known geometry of the solution space to perform allocation and learning without disrupting previously learned input-output mappings, but this is not a literal proposal for how biology addresses the continual learning problem. In particular, allocation and learning can completely avoid interference by modifying synaptic weights along the intended *η*-coordinates. However, in biology, plasticity must occur in the coordinate system of individual synaptic weights, and determining the required weight changes in *η*-coordinates requires knowing all past, current, and future input patterns (Eq. 6). There is some evidence that memory representations are fixed prior to memory encoding (69), so biology may learn with predetermined representations whose statistics make learning easier in *η*-coordinates. For instance, if the input patterns are all orthogonal to each other, then allocating and learning in *η*-coordinates merely requires synaptic plasticity based on new input patterns. While this scenario likely does not represent most learning, it illustrates what a brain must somehow accomplish to achieve its impressive continual learning capabilities (70). In less restrictive conditions, allocation could occur without disrupting previously stored memories by transiently changing the neuronal excitability, as discussed above. Nevertheless, the learning required to store new mappings and restore the original excitability would typically degrade older memories.

### Towards empirically accurate models of representational drift

Here we modeled representational drift for a fixed linear readout, and it is critical to model the solution space manifold under more biologically realistic scenarios. First, prior studies have suggested that readout weights also change during drift (19, 36). Our mathematical framework should thus be adapted to incorporate networks with plastic readout weights. Our preliminary analyses show that sparse solutions remain common when the representation and readout weight drift independently (SI Appendix A.2, Suppl. Fig. 2). Future work should account for changes coordinated across layers. Second, while this study modeled the readout as a linear projection, experimental evidence predominantly highlights non-linear stable subspaces that persist despite drift (20, 23–25). Extending the framework to incorporate non-linear readouts aligned with experimental observations is a promising direction for future work.

Biologically realistic models of representational drift also demand that parameter values be chosen to mimic experimental results. For instance, in this study’s proof-of-concept models, we classified both inactive and saturated neurons as disengaged because they both appeared frequently in the solution space and enhanced robustness to perturbations that posed challenges for gradient-based learning. Yet inactive neurons are far more prevalent than saturated ones in biology (71). Our model can mimic this outcome through parameter modifications (SI Appendix C.2, Suppl. Fig. 7). Moreover, we find that drift in these networks produces more biologically realistic single-cell dynamics, including intermittent firing and shifts in activity fields (36, 72). These *sparsely active* representations retain the robustness benefits highlighted for sparsely engaged ones, and also offer additional advantages, like energy efficiency (73–75) and robustness to readout changes.

### Further extensions to the theoretical framework

Future work should relax theoretical assumptions made for mathematical convenience, which may not be biologically correct. First, our framework assumes that the number of memories is smaller than the number of input neurons (*P* ≤*N*_*x*_), which enabled us to find a basis that separates out relevant weight combinations (32, 33). The existence of representational drift suggests that the brain may be highly redundant and operate below its memory capacity (11, 76), but this memory capacity can exceed the number of input neurons. Therefore, extending our framework to locally define the solution space manifold when the number of memories exceeds the number of neurons but remains below the network’s full capacity would be valuable. Second, drift may not be restricted to exact solutions but instead tolerate small non-zero errors in the readouts (15, 41, 48). It’s therefore important to generalize the framework to accommodate non-zero error solution spaces (32).

Extensions to the modeling framework could reveal higher-dimensional solution spaces that may be more robust. In this work, we observed that the solution space dimensionality was the same for densely-engaged and sparsely-engaged solutions (SI Appendix B). However, if certain regions of the solution were higher-dimensional, then this would provide a plethora of even more robust solutions. Here we observed such high-dimensional solution spaces in toy examples for specific values of the readout and activity thresholds (SI Appendix B.1, Suppl. Fig. 3), but they did not emerge in drift simulations (SI Appendix B.2). It’s possible that these solution space components are absent in the simulated examples. Higher-dimensional solution spaces may be more prevalent in extended models with dynamic neuronal excitability, where thresholds can be tuned (77, 78), or in models allowing non-zero readout errors, which have been shown to sometimes increase the dimensionality of solution spaces (35). It is also possible that higher-dimensional solution spaces exist but are hard to diffuse into. Developing analytical techniques to calculate the volume and dimensionality of solution spaces is thus crucial for determining whether drift can enhance robustness by utilizing higher-dimensional solution spaces.

### Model predictions and relation to experimental findings

In conjunction with neuroscience, our model makes several predictions. First, it predicts changes in engaged sparsity—or, in a more biologically relevant context (SI Appendix C.2, Suppl. Fig. 7), activity sparsity— proposing that representations initially become dense to facilitate learning, then sparsify during early drift, and maintain this sparsity during later drift. This aligns with findings in the CA1 hippocampus, a region known to have extensive drift (6, 8). In this region, activity in a novel environment increases through recruitment of additional active place cells, which is followed by sparsification with familiarization (79–82). Similarly, in the somatosensory cortex, another region exhibiting drift, neurons transiently increase activity during training before sparsifying (21, 60). While sparsification has been linked to learning (82) or behavioral changes (20), our model supports prior theoretical work (41) suggesting that it can arise due to solution space exploration. In contrast, regions with limited drift like CA3) don’t exhibit sparsity changes with familiarization (5, 83). Further supporting our findings, the fraction of active neurons remains stable during later drift(5, 7).

Second, our model predicts that representational drift rates can vary despite constant synaptic dynamics. Synaptic changes onto disengaged neurons do not affect representations unless thresholds are crossed. Therefore, sparsely engaged representations with many disengaged neurons should drift less than densely engaged ones. Supporting this, in V1 natural movies elicit denser representations that drift more, while gratings yield sparser more stable ones (84, 85). However, our model would require different bias terms for natural image representations and grating to generate different sparsity representations. During initial drift, we expect dense representations, so our model predicts greater drift, which is expected to slow down during later drift as representations sparsify (SI Appendix C.1, Suppl. Fig. 6). Further, drift rate may decrease as disengaged neurons move further into semi-constrained dimensions, making them less likely to rengage. This decline in the rate of representational changes aligns with observations of experiments (22, 72, 86, 87) and computational models (41). While the roles of learning and drift remain unclear, our model predicts that representational changes may continue to slow after learning concludes.

Third, our model predicts that representational drift results in large and variable input currents onto disengaged neurons. In networks with negative bias, sparsely active representations dominate, with inactive neurons receiving negative input currents well below the activity threshold (SI Appendix C.2 Suppl. Fig. 7). This contrasts with balanced network theory, which suggests near-threshold membrane potentials through finely tuned excitation and inhibition (88). While excitation and inhibition are balanced under anesthetization, inhibition dominates in states more relevant to memory representation, resulting in hyperpolarized neurons with membrane potential far below the spiking threshold (89). This phenomenon is seen across many brain regions that show extensive drift (90–92). Our model also predicts significant variability in subthreshold membrane potentials. Interestingly, high subthreshold potential variability is observed, but in a region with limited drift (93, 94). Further investigation into subthreshold potential variability across brain regions and its progression during learning and representational drift is crucial to rigorously test this model prediction.

## Materials and Methods

We established a task framework and neural network architecture to study representational drift. The input matrix *X*, comprising the activity of *N*_*X*_ = 15 input neurons across *P* = 8 input conditions, was drawn from *X* ∼ 𝒩 (0, 1). The representation-layer consisted of *N*_*Y*_ = 12 neurons and its activity *Y* was used to generate an *N*_*Z*_ = 1-dimensional readout, through a fixed readout weight matrix *W* ∼ 𝒩 (0, 10). We used a clipped-threshold-linear activation function with an activity threshold of 0 and a saturation threshold of *α* = 5, unless specified otherwise. The desired readout is specified by a teacher network, which processes the input through a weight matrix *T*, applies the same clipped-threshold-linear activation function, and performs a one-dimensional linear projection via the same weights *W*. Thus, the desired readouts are given by *W* Φ(*TX* + *B*), while the readout generated by the network is given by *W* Φ(*UX* + *B*), where *B* is the bias term, which was set to a zero matrix unless stated otherwise.

To simulate drift as an exploration of the solution space, we first reached the solution space through learning. The weights *U* were initialized as *U* (0, 0.1) unless specified otherwise. We allocated the weight combinations relevant to the *P* input conditions within the center of the engaged regime by setting 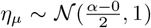, where 0 ≤ *µ < P*. Subsequently, we updated the corresponding weights *U* using gradient descent (Adam optimizer in PyTorch, with a learning rate of 0.001) to minimize the Mean Squared Error (MSE) loss between the network readouts and the desired readouts generated by the teacher network across the *P* input conditions. The readout weights *W* and the bias term *B* = 0 were kept fixed unless specified otherwise. When adding the *µ*th memory in the continual learning setting with gradient descent directly in *U* (Fig. 7F-I), we only allocated *η*_*µ*_ and updated the corresponding weights *U* to minimize the loss only for the new memory *µ* using Eq. 9. Similarly, for continual learning with gradient descent in *η*-coordinates (Fig. 7A-E), we allocated *η*_*µ*_ and then updated it to reduce the loss for the *µ*th input condition using Eq. 10.

We would like to acknowledge the use of OpenAI’s ChatGPT for assisting with language and grammar refinement and Adobe Illustrator’s generative AI for help creating Fig. 1A.

## Acknowledgements

The authors thank Tim O’Leary, Nelson Spruston, Jim Knierim, and members of the Fitzgerald Lab for useful discussions. This work was supported by the Howard Hughes Medical Institute. JEF acknowledges institutional support to the National Institute for Theory and Mathematics in Biology from the National Science Foundation (grant number DMS-2235451) and the Simons Foundation (grant number MPTMPS-00005320).

## SI Appendix

**Supplementary Figure 1.**
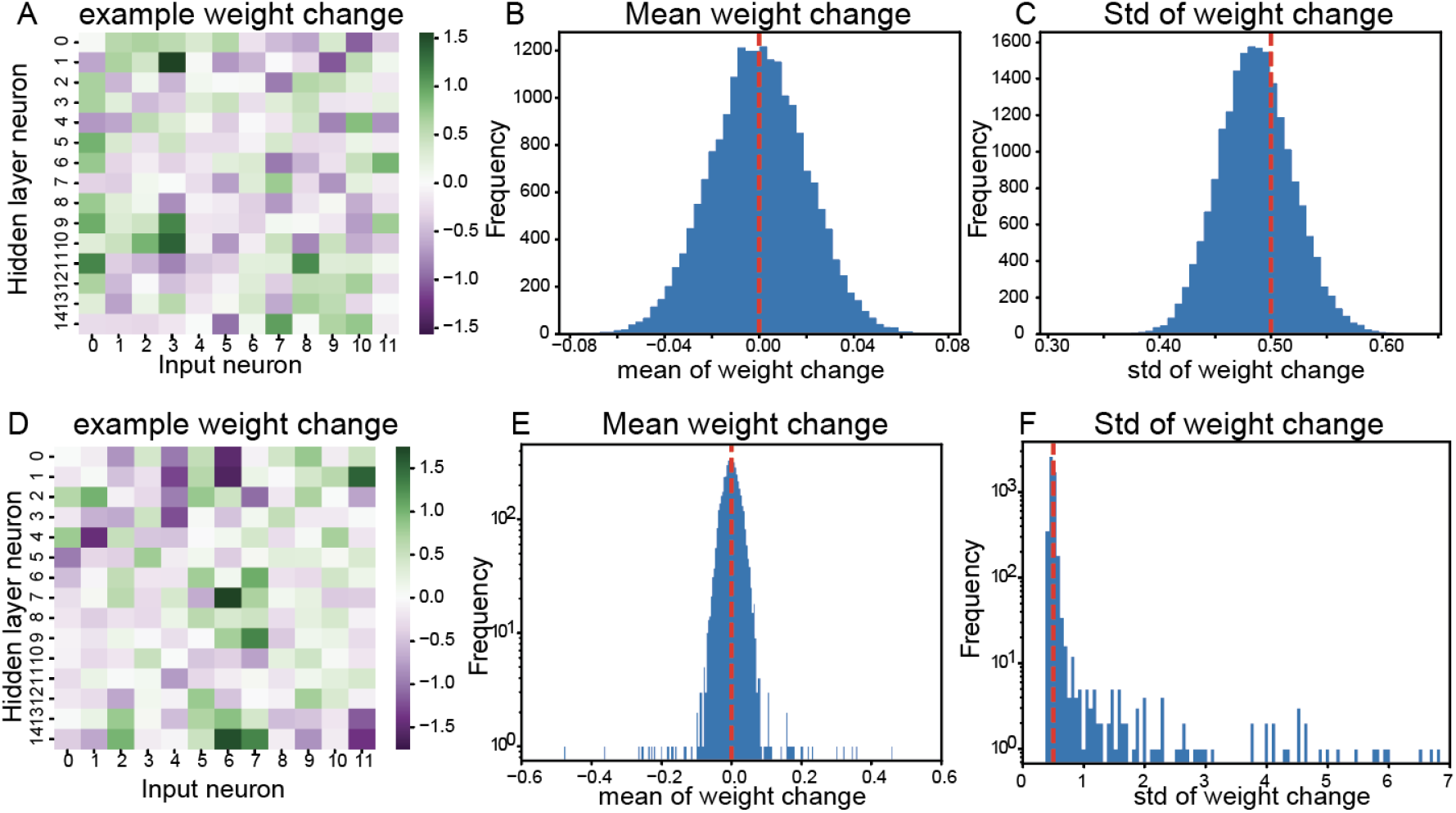
Characterization of synaptic weight changes during drift. (A-C) Drift without any weight norm bound. (A) An example *dU* resulting from uncorrelated *η* change. (B) The distribution of mean weight changes, with red dotted line indicating the expected mean of 0 (C) The distribution of the standard deviation of weight changes, which is slightly smaller than the expected standard deviation (*σ* = 0.5, marked by the red dotted line) possibly due to rescaling of changes done to avoid threshold crossing. (D-F) Drift with weight norm bound set to its initial value for each neuron. (D) Example weight change *dU*. (E) Distribution of mean weight changes is still centered around 0. However, some weight changes exhibit means further away from 0 due to additional changes made to ensure weight norm constraint. (F) The distribution of the standard deviation of weight changes remains close to the expected *σ* = 0.5, although there are instances of larger weight changes, made to adhere to weight norm constraints.

### A. Model variants

#### A.1. Generalizing to other non-linear continuous activation functions

In this section, we generalize the proposed solution space manifold setup to accommodate a broader class of non-linear activation functions. Consider a network where the representation-layer activity is given by *Y*_*i,µ*_ = Ψ (Σ_*j*_ *U*_*i,j*_*X*_*j,µ*_), where *Y*_*i,µ*_ is the activity of the *i*^*th*^ neuron under input condition *µ, U*_*i,j*_ is the synaptic weight from input neuron *j* to representation-layer neuron *i, X*_*j,µ*_ is the input activity of neuron *j* under input condition *µ*, and Ψ represents a continuous activation function defined as

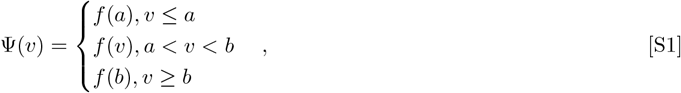

where *f* (*v*) is a strictly increasing function, ie. if *a* ≤ *v*_1_ *< v*_2_ ≤ *b*, then *f* (*v*_1_) *< f* (*v*_2_). We define another function Ω

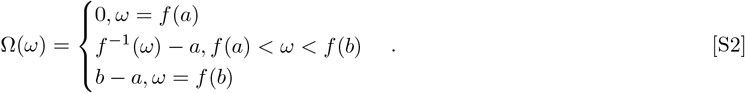

Then

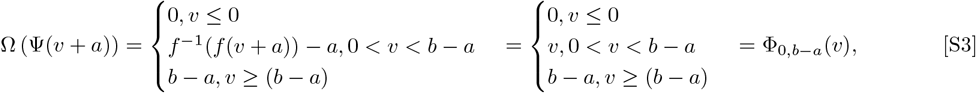

where Φ_0,*b*−*a*_(*v*) is a clipped-threshold linear activation function with an activity threshold of 0 and a saturation threshold of *b* − *a*. If we define 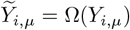, then the original network with Ψ activation function becomes equivalent to a network with clipped-threshold linear activation function Φ_0,*b*−*a*_ with an activation threshold of 0, a saturation threshold of *b* − *a* and a baseline activity or bias of −*a*

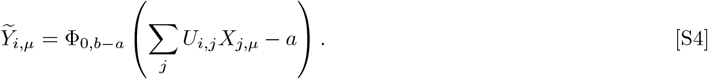

#### A.2. Generalizing to dynamic readout weights

In this paper, we modeled representational drift by generating different representations that satisfy a particular input-output mapping with fixed abstract readout weights. Here, we extend our approach to a scenario where both the input synaptic weights *U* and the abstract readout weights *W* are subject to instability (Suppl. Fig. 2A). We do so by alternating between two steps: (1) keeping *W* fixed and identifying changes in *U* that preserve the downstream readout, and (2) keeping *U* (and subsequently *Y*) fixed and determining changes in *W* that maintain a fixed readout (specifically, changes in the null space of *Y*).

Previously, instead of making changes directly in *U*, we made changes in a transformed version of *U* ie. *η*. Similarly, we make changes in a basis transformed version of *W*, ie. *ζ* We consider the case where the number of neurons in the hidden layer is greater than the number of stimuli stored ie. *N*_*Y*_ *> P*,

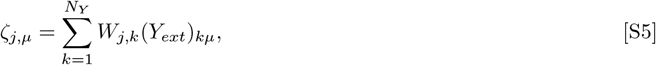

where *Y*_*ext*_ is computed from *Y* by stacking *N*_*Y*_ − *P* columns in the null space of *Y* to it. Then we make changes *δζ* as follows:

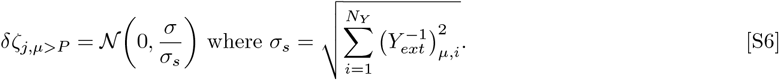

Similar to the Weight norm bound set to the hidden layer neurons, for each dimension of the readout *k*, we impose a bound on the readout weights such that

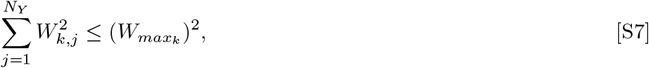

where *W*_*max*_ is the maximum weight norm allowed onto the *k*^*th*^ readout dimension. When a proposed *δζ* causes the weights onto readout *k* to exceed its limit 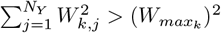, then we scale the unconstrained dimensions of *δζ* such that the weight norm is set to the bound ie. 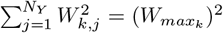. Thus, we modified synaptic weights *U* and readout weights *W* (Suppl Fig. 2B-D), while preserving the relevant readouts *Z* (Suppl. Fig. 2E).

While this method allowed us to generate representational changes beyond the null space of the readout weights *W*, it may not comprehensively sample from the full combined solution space of (*U, W*). To illustrate, consider a simplified example where a single input neuron projects with weight *u* to a hidden layer neuron, which further projects with weight *w* to a readout neuron. The readout is given by *z* = *w*Φ(*ux*), where Φ represents a clipped-threshold linear activation function and *x* is the input. The task entails maintaining a readout *z* = 2 for an input *x* = 1, which simplifies to the constraint *w*Φ(*u*) = 2. When the neuron is engaged, this further reduces to *wu* = 2, indicating that there are many solutions of synaptic and readout weight combinations, and the system should be able to diffuse in the solution space (Suppl. Fig. 2F). Now, consider starting from an initial solution of (*u, w*) = (1, 2). First, we fix *w* = 2 and attempt to change *u* to maintain the readout. However, this leads to 2*u* = 2, fixing *u* at 1. With only one input neuron (*N*_*X*_ = 1) and one stimulus to store (*P* = 1), there are no unconstrained dimensions. Furthermore, since there are no disengaged neurons, there are no semi-constrained dimensions. The number of hidden-layer neurons (*N*_*Y*_ = 1) matches the readout dimensionality (*N*_*Z*_ = 1), meaning the hidden-layer activity cannot change without altering the readout. Consequently, *u* cannot be changed. Next, we attempt to modify *w* while keeping *u* fixed, but this also lacks flexibility, as it requires *w* = 2. Thus, the system becomes stuck in the solution (*u, w*) = (1, 2). However, this solution is surrounded by solutions. Exploring these requires simultaneous changes in both *u* and *w*, which this method cannot achieve. Thus, joint modification of *u* and *w* is necessary to access certain solutions, highlighting a limitation of this approach.

Robustness to synaptic weight perturbations persists in this setup, but inactive and saturated neurons exhibit opposing effects when considering readout weight changes. Representational drift generated using this method, alternating *U* and *W* changes still led to sparsely engaged solutions (Suppl. Fig. 2G). As before, these solutions showed enhanced robustness to input weight perturbations (Suppl. Fig. 2H). Since this setup allows changes to both input synaptic weights and abstract readout weights, we also considered robustness to *W* readout weight perturbations. The error in readout *Z* due to a perturbation in *W* can be quantified as *dWY*, indicating that the error is proportional to the activity level of representation-layer neurons. Neurons with no activity (i.e., inactive neurons *Y*_*j,µ*_ = 0) do not contribute to the error, whereas neurons that have reached their maximum activity level (i.e., saturated neurons *Y*_*j,µ*_ = *α*) contribute the most to the error. Drift led to an increase in both inactive and saturated neurons, resulting in post-drift solutions exhibiting similar robustness to readout weight perturbations compared to pre-drift solutions (Suppl. Fig. 2I). Although both engaged and saturated neurons enhanced the robustness of the model to perturbations in the input synaptic weight *U*, they have opposing effects on the robustness to changes in *W*. Therefore, when the readout weights themselves are subject to change, sparsely active solutions may offer greater robustness compared to sparsely engaged ones.

**Supplementary Figure 2.**
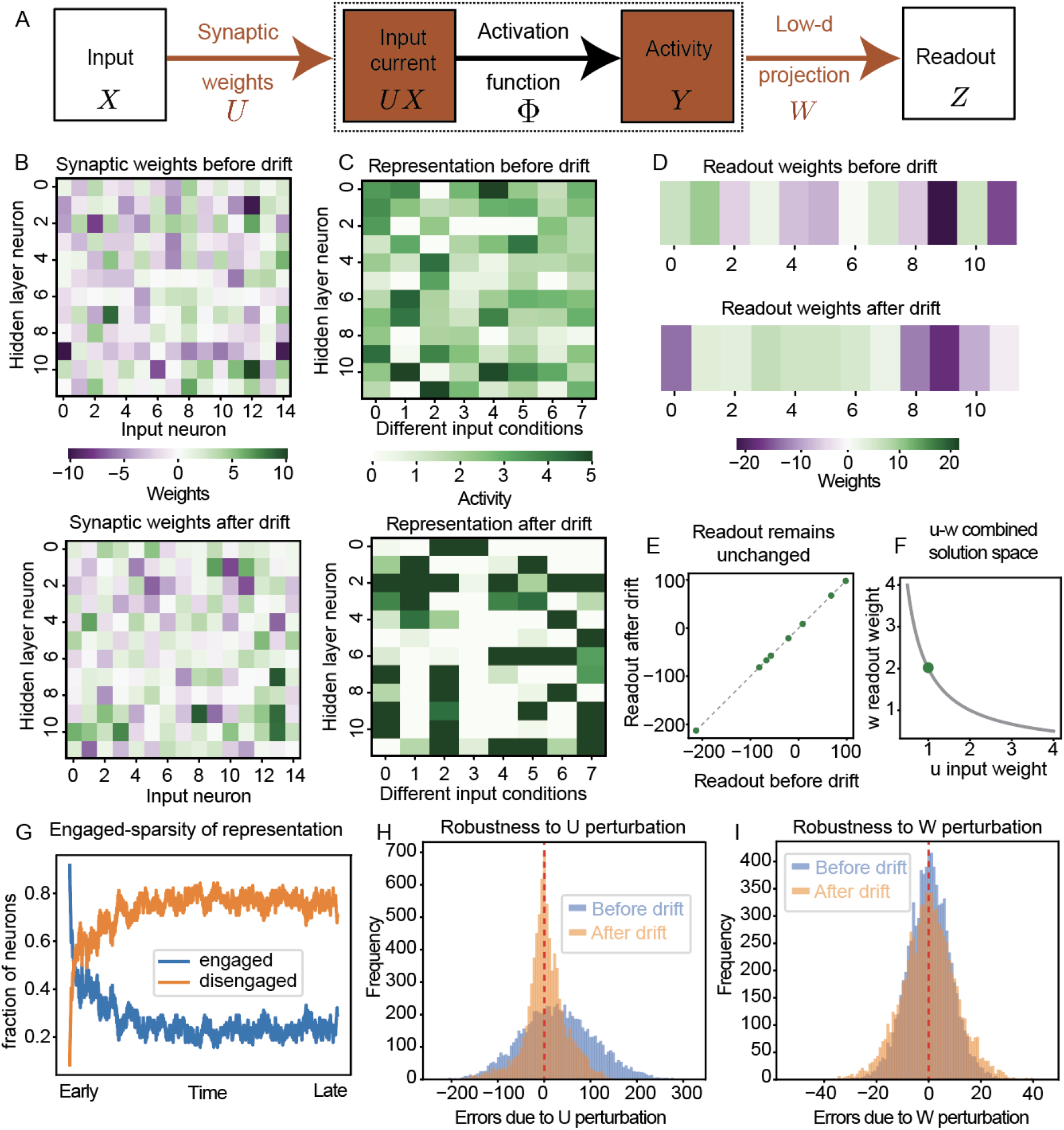
Generalizing to dynamic readout weights (A) Network framework with dynamic readout weights. (B) Synaptic weights before and after drift demonstrating synaptic turnover. (C) Neural activity under different input conditions before and after simulating drift. (D) Readout weights before and after drift. (E) Readout for each input condition remains the same before and after drift. (F) Toy example solution space to illustrate the need for coordinated changes in input and readout weights. (G) Fraction of engaged and disengaged (inactive and saturated) neurons over time during drift showing drift favors sparsely engaged representations. (H) Histogram of signed errors in readout due to U perturbations *N* (0, 1) showing greater robustness post-drift. (I) Histogram of signed errors in readout due to W perturbations *N* (0, 1) showing sparsely engaged representations don’t enhance robustness to W perturbations.

### B. Understanding the geometry and statistical properties of solution spaces

#### B.1. Illustrative toy problem

To illustrate why robust solutions might emerge more naturally through drift than through learning, we’ll analyze the solution space and error gradients for a simple toy model. The toy model consists of a two-neuron representation *y* (Suppl. Fig. 3A), which receives input from a single sensory neuron *x* and influences a 1-dimensional linear readout *z*. We consider a task where the system needs to reproduce a single mapping from input *x* to output *z*.

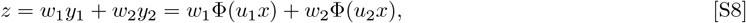

where *w*_1_ and *w*_2_ parameterize the readout, *x* is the input, *y*_1_ and *y*_2_ are the activities of the two neurons, and Φ (Suppl. Fig. 3B) is the clipped-threshold-linear activation function given by

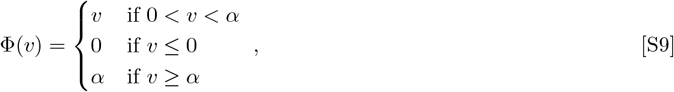

where the activity threshold is 0 and saturation threshold is *α*. Between 0 and *α*, changes in input currents lead to changes in neural activity, so we denote neurons in this regime “engaged”. Neurons with input currents below the activity threshold 0 are inactive and those with input currents exceeding the saturation threshold *α* are saturated. Both inactive and saturated neurons exhibit activity invariance to small input current changes, thus we classify them as “disengaged”. In the examples that follow, we set *α* = 5, *x* = 1, (*w*_1_, *w*_2_) = (1, 1), and analyze the solution space of (*u*_1_, *u*_2_) for different values of readout *z*.

Synaptic weight solutions can vary in statistical properties, such as the sparsity of the representations they produce, with some level of sparsity being more prevalent than others. Suppose the required readout is *z* = 4, then the representation must satisfy *y*_1_ + *y*_2_ = 4. When both neurons are in the engaged regime (i.e., 0 *< y*_1_ *<* 5 and 0 *< y*_2_ *<* 5), we get densely engaged solutions satisfying *u*_1_ + *u*_2_ = 4 (Suppl. Fig. 3C, blue line). Since flexibility along this dimension changes the neural representations, we term this as “drift dimension”. At (*u*_1_, *u*_2_) = (0, 4), the first neuron becomes inactive, so its input current only needs to satisfy *u*_1_ ≤ 0, giving rise to sparsely engaged solutions with one inactive and one engaged neuron (Suppl. Fig. 3C orange line) Since the inequality constrains the solution space to part of a dimension, we denote this as “semi-constrained” dimension of flexibility. We also find another semi-constrained dimension *u*_2_ ≤ 0 when *u*_1_ = 4, which also gives rise to sparsely engaged solutions. As the weight norm bound increases, larger input currents are allowed, expanding the volume of sparsely engaged solution spaces with an inactive neuron (Suppl. Fig. 3C,D). In contrast, the size of the densely engaged solution space remains unchanged. For sufficiently large weight norms, the volume of sparsely engaged solution space surpasses that of densely engaged solution space, causing diffusion to favor these sparsely engaged representations.

Sparsely engaged representations are robust to noise. When a random perturbation is applied to synaptic weights (*u*_1_, *u*_2_), it affects the input currents that neurons receive, which in turn affects the activity of engaged neurons and the readout. For inactive neurons, the activity remains at 0 if the perturbation is small enough that the input current doesn’t cross the activity threshold, so these neurons don’t contribute to downstream error. Therefore, a weight perturbation of the same magnitude leads to a larger error when starting from a densely engaged solution than when starting from a sparsely engaged solution (Suppl. Fig. 3E,F). Saturated neurons also contribute to such increased robustness of sparsely engaged representation solutions. Small weight perturbations that do not cause the input current to fall below the saturation threshold do not affect the activity of a saturated neuron. Indeed, when the required readout is *z* = 6, we find that sparsely engaged solutions involving saturated neurons (*u*_1_ ≥ 5, *u*_2_ = 1 or *u*_1_ = 1, *u*_2_ ≥ 5)(Suppl. Fig. 3G, orange lines) are more robust to weight perturbations (Suppl. Fig. 3H,I) than densely engaged solutions satisfying *u*_1_ + *u*_2_ = 6 (Suppl. Fig. 3G).

**Supplementary Figure 3.**
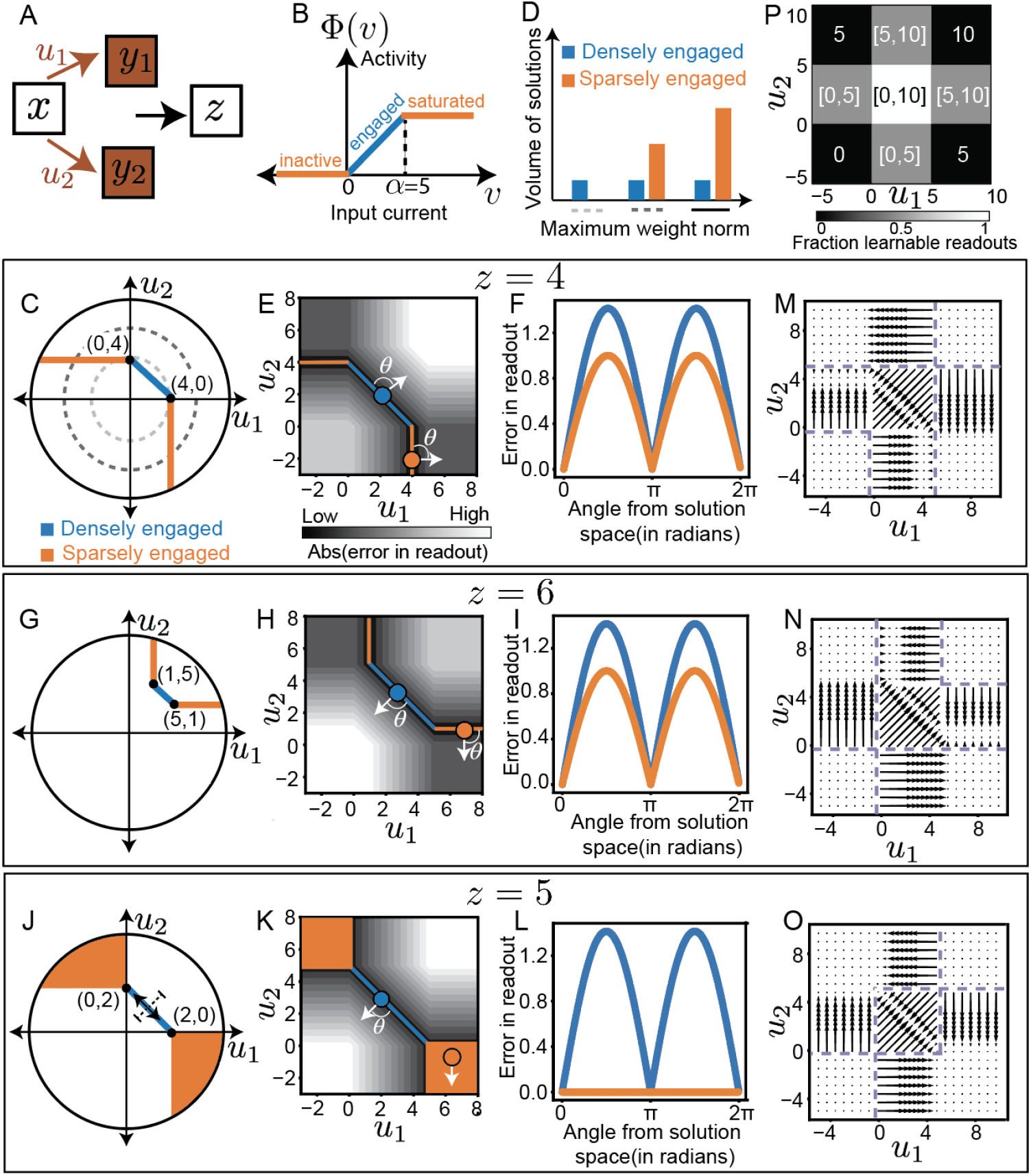
A toy example illustrating the prevalence, robustness, and computational challenges of learning sparsely engaged solutions. (A) Network architecture for a toy example whose task is to generate a fixed readout, *z*, from two-dimensional neural activity, (*y*_1_, *y*_2_), generated from a single sensory input, *x*, via weights, (*u*_1_, *u*_2_). (B) Activation function Φ indicating the regime for engaged, inactive, and saturated neurons. Subsequent panels examine solutions for different readout values: *z* = 4 (C-F); *z* = 6 (G-I); and *z* = 5 (J-L). (C) Synaptic weight solution space (*u*_1_, *u*_2_) for *z* = 4, with sparsely engaged solutions in orange and densely engaged solutions in blue. Dotted lines indicate different weight norms. (D) The volume of sparsely engaged solution space increases with increase in the weight norm bound. Line styles along the x-axis indicate the weight norm bounds shown in (C). (E) Error in readout *z* for various (*u*_1_, *u*_2_) and a schematic demonstrating weight perturbations (white arrow) from sparse (orange) and densely engaged (blue) solutions. (F) Weight perturbations from sparsely engaged solution (*u*_1_, *u*_2_) = (4, − 2) produces lower readout error than perturbations from a densely engaged solution, (*u*_1_, *u*_2_) = (2, 2) across all perturbation angles *θ*. (G,H,I) Same as (C,E,F) but with the desired readout *z* = 6, showing that sparsely engaged solutions with a saturated neuron (orange) have enhanced robustness similar to sparsely engaged solutions with an inactive neuron. Perturbations are from a sparsely engaged solution with one saturated neuron, (*u*_1_, *u*_2_) = (1, 7) (orange dot), or a densely engaged solution, (*u*_1_, *u*_2_) = (3, 3) (blue dot). (J,K,L) Same as (C,E,F) but with the desired readout *z* = 5 showing a 2-dimensional space of maximally sparsely engaged solutions in orange and a 1-dimensional space of densely engaged solutions in blue. Perturbations are from a maximally sparsely engaged solution with a saturated neuron and an inactive neuron (*u*_1_, *u*_2_) = (7, − 2) (orange dot), and from a densely engaged solution, (*u*_1_, *u*_2_) = (2.5, 2.5) (blue dot). (M, N, O) Error gradient flow fields plots for desired readout *z* = 4 (M), *z* = 6 (N), and *z* = 5 (O). Arrows indicate the direction in which errors decrease, and their length corresponds to the gradient magnitude. Dotted lines delineate the basin of attraction reached from initialization at that location. (P) For each of the 9 regions with different fraction of engaged, inactive, and saturated neurons, the value/interval of *z* readout for which the solution space can be reached by following the gradient. Color indicates the fraction of learnable readouts *z*.

Different dimensional solution spaces may coexist for particular mappings. When the required readout is *z* = 6, consider a system diffusing within the densely engaged solution space, satisfying *u*_1_ + *u*_2_ = 6. When the system reaches (*u*_1_ = 5, *u*_2_ = 1), the first neuron becomes saturated, introducing a semi-constrained dimension (*u*_1_ ≥ 5). However, this flexibility is offset by a loss of drift dimension flexibility because the second neuron must now satisfy *u*_2_ = 1 with no flexibility (Suppl. Fig. 3G). Since a drift dimension converts into a semi-constrained dimension, the overall dimensionality of the solution space remains unchanged before and after the first neuron becomes saturated. Now, consider a different required readout of *z* = 5. Here, two types of solutions emerge: (1) Densely engaged solutions with both engaged neurons satisfying *u*_1_ + *u*_2_ = 5 (2) Maximally sparsely engaged solutions with one inactive and one saturated neuron, satisfying two inequality constraints (ex. *u*_1_ ≥ 5, *u*_2_ ≤ 0) (Suppl. Fig. 3J). Suppose the system explores the densely engaged solution space *u*_1_ + *u*_2_ = 5 and reaches (*u*_1_ = 5, *u*_2_ = 0). At this point, the first neuron becomes saturated and the second becomes inactive. This adds two semi-constrained dimensions (*u*_1_ ≥ 5 and *u*_2_ ≤ 0) while losing one drift dimension flexibility, increasing the dimensionality of the solution space to two. Thus, a two-dimensional maximally sparsely engaged solution space coexists with a one-dimensional densely engaged solution space.

A higher-dimensional solution space implies greater prevalence and robustness. Since maximally sparsely engaged solutions form a two-dimensional solution space, they are infinitely more numerous than densely engaged solutions, which are confined to a one-dimensional space. Furthermore, small weight perturbations in the maximally sparse solution do not alter the state: the inactive neuron remains inactive, the saturated neuron remains saturated, and both the activity and readout remain unchanged. This makes maximally sparse solutions not only more prevalent but also maximally robust (Suppl. Fig. 3K,L). Once the system reaches this robust, higher-dimensional solution space, transitioning back to the densely engaged solution space becomes highly improbable. To do so, the system would need to shift from exploring a two-dimensional region (*u*_1_ ≥ 5 and u_2_ ≤ 0) to a single point (*u*_1_ = 0, *u*_2_ = 5) a transition that is unlikely to occur through diffusion, thereby preserving this robust representation.

These toy examples have demonstrated that sparsely engaged representations are common, implying drift will favor these solutions. Interestingly, we will see that despite being more common, these solutions are challenging to learn and can hinder subsequent learning. To do so, we will examine regions of the synaptic parameter space that facilitate learning and the types of solutions that emerge when starting from these learning-conducive regimes. Consider a scenario where the goal is to learn a mapping from an input *x* to a desired readout *z*. A random weight configuration likely produces a readout 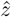 that differs from the desired readout *z*, resulting in a squared readout error of 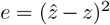. To learn the mapping, the weights need to change to minimize this readout error. Learning the mapping requires adjusting the weights to minimize this error. This process is often modeled using gradient descent, where weights are updated in the direction that most effectively reduces the error—specifically, the direction opposite to the gradient of the error function 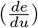. Since gradient information is essential for learning, we will analyze the gradients at different weight configurations for different required readouts *z* while setting a fixed input *x* = 1, to identify regions of the parameter space that are most conducive to learning.

Maximally sparsely engaged initializations are unsuitable for learning because they provide no gradients. If the initial weights cause both neurons to become disengaged, small changes in input currents have no effect on neuronal activity, the readout, and the readout error. As a result, the error function is locally flat and lacks gradients (Suppl. Fig. 3M,N,O). This prevents the weights from updating, leaving the system stuck in a local optimum regardless of the required readout. These initializations are only effective if the current readout 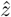 already matches the desired readout *z* (Suppl. Fig. 3O). For instance, when the required readout is *z* = 5 and if we initialize at (*u*_1_, *u*_2_) = (−2, 7) which produces a maximally sparsely engaged representation (0, 5), the network would already produce the correct readout 5, requiring no learning.

Partially sparsely engaged initializations face similar but less severe challenges compared to maximally sparse ones. Consider an initialization where the first neuron is engaged and the second is inactive, such as (*u*_1_, *u*_2_) = (1, −4). If the desired readout *z* permits a solution where the second neuron remains inactive (e.g., for *z* = 4, solutions like (*u*_1_, *u*_2_) = (4, −4) exist), the system can leverage gradients from the engaged neuron to converge to these sparsely engaged solutions (Suppl. Fig. 3M). However, if no such solution exists with the second neuron being inactive (e.g., for *z* = 6), the system initially reduces the error by increasing the activity of the first neuron. However, once the first neuron saturates, the system becomes trapped in a local optimum where both neurons are disengaged. For instance, starting from (*u*_1_, *u*_2_) = (4, − 4), the system may increase *u*_1_ to reach (*u*_1_, *u*_2_) = (6, − 4). Here, the network generates neural activity of (*y*_1_, *y*_2_) = (5, 0) and a readout error of 1. Since both neurons are disengaged, there are no gradients and the system will remains stuck (Suppl. Fig. 3N). Thus, while partially sparse initializations can succeed, they frequently get stuck in local optima (Suppl. Fig. 3P), making them less conducive for effective learning.

Densely engaged initializations are the most conducive to learning but tend to yield non-robust solutions. For example, consider an initialization at (*u*_1_, *u*_2_) = (1, 1), where both neurons are engaged. For any readout within the feasible range (0 ≤*z* ≤ 10), this and other densely engaged initializations can reliably find a solution (Suppl. Fig. 3 M-P). However, while densely engaged initializations succeed in finding solutions, they often converge to non-robust, densely engaged solutions that are closer in parameter space, even when robust, sparsely engaged solutions are more prevalent. For instance, if the initialization is (*u*_1_, *u*_2_) = (1, 1) and the desired readout is *z* = 4, gradient-descent learning would follow the trajectory of maximal error reduction along the line *u*_1_ = *u*_2_ ultimately reaching the solution space at (*u*_1_, *u*_2_) = (2, 2) (Suppl. Fig. 3M, left). This solution, however, is densely engaged and non-robust. Consequently, a mechanism like drift may be advantageous in guiding the system from densely engaged, non-robust solutions that are easy to learn to sparsely engaged, robust solutions that are more common.

To summarize, these toy examples suggest that given a sufficiently large weight norm, sparsely engaged, noise-robust solutions are more common and thus preferred by drift. Moreover, these robust solutions are harder to reach through learning when starting from learning-conducive regions, suggesting drift following learning may help find these solutions. The next few subsections will investigate the prevalence and robustness of drifting representations.

#### B.2. The sparsity of solutions explored is influenced by the input current exploration ranges of engaged and disengaged neurons

Here, we seek to better understand the relationship between solutions explored by drift and their prevalence. As weight norm bounds increased, we observed that drift-explored representations became more sparsely engaged and robust. We hypothesize this occurs because larger weight norm bounds enable larger input currents, expanding exploration ranges for inactive and saturated neurons, and consequently increasing the prevalence of sparsely engaged solutions. To validate this hypothesis, we directly impose input current limits instead of weight norm bounds and compare the sparsity of drift-explored representations to their expected solution prevalence. To simplify prevalence computation, we approximate solution frequency with the overall representation frequency at different sparsity levels, irrespective of solution status.

**Supplementary Figure 4.**
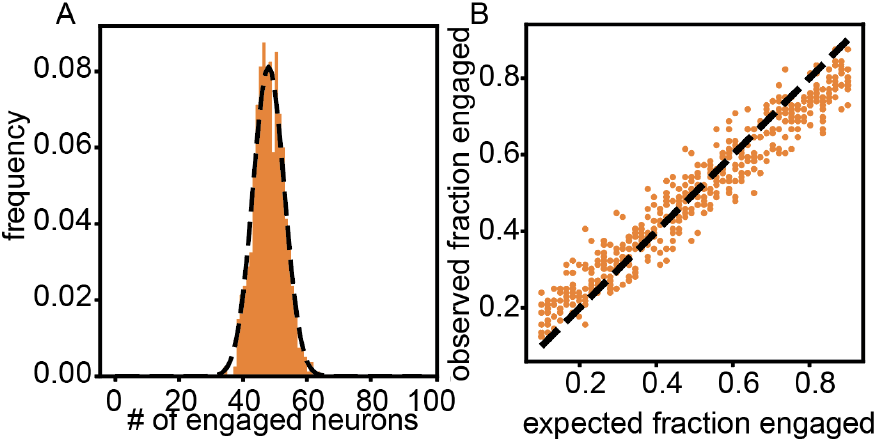
Understanding the sparsity of solutions achieved by drift. (A) The orange histogram shows the frequency of representations having a certain number of engaged neurons *E* during drift with parameters *c*_1_ = −2.5 and *c*_2_ = 7.5. To avoid correlated data, we sampled only once every 500 steps and began sampling after 5000 timesteps to ensure equilibration. The dotted line represents the theoretical prediction for the probability of representations with different number of engaged neurons given by *B*(*N*_*y*_ *P* = 96, *p*_*E*_ = 0.5), demonstrating a good match between the theoretical sparsity prediction and the sparsity of representations explored by drift. (B) For 50 simulations of drift with different *c*_1_ and *c*_2_ values chosen to explore a wide range of engaged neuron fractions (0.1 ≤ *p*_*E*_ ≤ 0.9), the x-axis shows the expected fraction of engaged neurons, while the y-axis shows the observed fraction of engaged neurons. Sampling was performed every 500 steps for each simulation to avoid correlated data.

We analyze the same model configuration, consisting of *N*_*x*_ input neurons, *N*_*y*_ neurons in the representation layer, and *N*_*z*_ abstract readouts. The task involves maintaining a mapping between input *X*_*µ*_ and corresponding readout *Z*_*µ*_ for *µ* = 1, ‥, *P* mappings. The weights *U* onto representation-layer *Y* are plastic and the readout is defined by a fixed matrix *W*. Consider limiting the input current range to (*c*_1_, *c*_2_), where *c*_1_ ≤0 and *c*_2_ ≥ *α*. Under these conditions, a neuron is engaged with probability 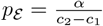. Consequently, the number of engaged neurons, 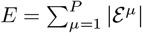, is binomially distributed as

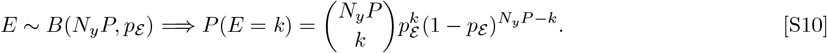

For instance, if *c*_1_ = − 2.5 and *c*_2_ = 7.5 then the probability of being engaged and disengaged are the same at 0.5. Here, we observed an excellent match between this theoretical distribution from Eq. S10 and the fraction of time points that drift spent in representations with varying numbers of engaged neurons (Suppl. Fig. 4A). By systematically varying *c*_1_ and *c*_2_ over the ranges −22.5 ≤*c*_1_ ≤−0.27 and −27.5 ≤*c*_2_ ≤ 5.27, we modulated drift to favor different sparsity levels from 0.1 to 0.9 (Suppl. Fig. 4B). The expected fraction of engaged neurons, *p*_E_, closely matched the observed mean fraction during drift, with minor discrepancies at extremely low or high sparsity. These findings support that sparsely engaged representations are favored by drift due to the inherent prevalence of these representations when input currents are permitted to be large.

While sparsely engaged solutions might have been favored due to higher-dimensionality of their solution spaces, in our simulations this is not the driving factor. Let there be *N*_*z*_ + *n*_*ℰ*_ engaged neurons for a particular input condition *µ*, where *N*_*z*_ is the dimensionality of the required readout. Since the number of engaged neurons is *n*_*ℰ*_ greater than the readout dimensionality, the engaged submatrix *W* ^*µ*^ has a *n*_*ℰ*_ -dimensional null space, implying there are *n*_*ℰ*_ drift dimensions for *µ*. During solution space exploration, if *n*_*ℰ*_ + 1 of those engaged neurons simultaneously become disengaged then *n*_*ℰ*_ + 1 semi-constrained dimensions are added. However, there were only *n*_*ℰ*_ drift dimensions for *µ* before, so only *n*_*ℰ*_ drift dimensions can be lost. Thus, the new solution space would become larger by one dimension compared to the previous solution space. In our drift simulations, for *N*_*z*_ = 1, this would require zero engaged neurons for an input condition. However, we observed no cases where the number of engaged neurons dropped to zero, suggesting that increased dimensionality of solution space was not responsible for exploration of sparsely engaged solutions.

Higher-dimensional solution spaces not being explored in our simulations raises intriguing questions about their feasibility and accessibility. As illustrated in the toy example (Suppl. Fig. 3), such higher-dimensional solution spaces arise only under highly specific conditions involving precise values of the readout, readout weights, and activity thresholds. It is plausible that, similar to some other toy examples explored with required readouts of *z* = 4, 6 the input-output mappings in our simulations may not have had such higher-dimensional solution spaces. Such higher-dimensional solution spaces may become more feasible if we allow non-zero readout errors, changes in readout weights, and dynamic activity, and saturation thresholds. Alternatively, such higher-dimensional solutions may indeed exist but could be inherently more challenging to reach with our current methods or through diffusion-driven exploration in general. Future work could focus on developing methods to access these maximally sparsely engaged and higher-dimensional solution spaces, potentially offering exceptional robustness and stability.

#### B.3. Robustness of sparsely engaged solutions stems from stable activity of disengaged neurons despite perturbations to input-current

Here, we delve deeper into the relationship between robustness and representational sparsity. Consider applying a weight perturbation *dU*, then the new downstream readout for the *µ*^*th*^ stimulus becomes 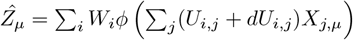. The readout error for the *µ*th stimulus 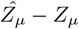 is given by

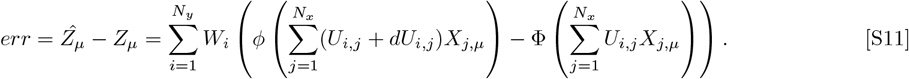

Assume *dU* is sufficiently small such that none of the neurons cross thresholds. Then for *i* ∈*/ ℰ*^*µ*^ (ie. disengaged neurons), we have 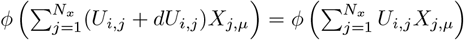, so these neuron-stimuli (*i, µ*) pairs don’t contribute to the readout error. Since we assume engaged neurons remain engaged after perturbation, we can remove the non-linearity to get

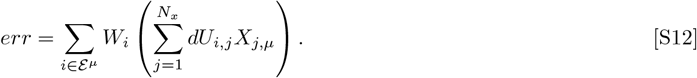

If *β* is the fraction of engaged neurons for input condition *µ* then *I* ∈ ℰ^*µ*^ only sums over *N*_*y*_*β* neurons. If *W, dU*, and *X* are uncorrelated random variables with standard deviations *σ*_*W*_, *σ*_*U*_, and *σ*_*X*_ respectively, then the standard deviation of the error due to perturbation is given by

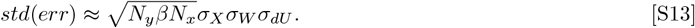

Drift promotes sparsely engaged solutions, characterized by fewer engaged neuron-stimulus pairs (lower *β*), leading to a reduced standard deviation of errors. By exploring solutions with varying sparsity *β* through limiting input currents during drift, we found a strong correlation between the ratio of standard deviation of errors before and after drift and the ratio of square root of the fraction of engaged neurons before and after drift, as predicted by Eq. S13 (Suppl. Fig. 5 A). This suggests that the robustness of post-drift solutions arises from sparsely engaged neurons maintaining their activity levels despite weight perturbations.

In our previously analysis, we assumed that the perturbation *dU* doesn’t lead to threshold crossing, although this assumption may not always hold. If the perturbation is sufficiently large or the input current is close to the thresholds, then weight perturbations may cause neurons to cross thresholds. When an engaged neuron crosses thresholds to become disengaged, the weight change onto that neuron till the threshold crossing contributes to an error, while the remaining perturbation onto that neuron doesn’t alter its activity, and thus doesn’t contribute to errors. Consider the previously described toy model in B.1 with a required readout *z* = 6 and a particular weight solution of (*u*_1_, *u*_2_) = (1.3, 4.7) (Suppl. Fig. 5 B). We find that perturbations from this solution, towards threshold crossing produce smaller positive errors than expected for a network without threshold non-linearity. Further, at threshold crossing, the solution space dimensions change, in this case from (*u*_1_ + *u*_2_ = 5) to (*u*_2_ *>*= 5), so a perturbation along the engaged solution space (*u*_1_ + *u*_2_ = 5) at angle *π* ends up producing negative errors. Consequently, the error due to perturbations are not symmetric around 0, in this case producing fewer positive errors than negative errors (Suppl. Fig. 5C). Which in turn manifests as a bias in the error, producing asymmetric error histograms (Suppl. Fig. 5D).

Solutions before drift show more bias in errors due to perturbation than solutions after drift. Crossing of the threshold becomes more likely when the input current is close to one of the thresholds, which in turn occurs more often when input drive exploration range is small. In our simulations where weight norms are allowed to be sufficiently large and no constraints are imposed on the input currents, disengaged neurons have a very large exploration range, while engaged neurons always have a small exploration range (0, *α*). As a result, engaged neurons are more likely to cross the threshold and become disengaged than disengaged neurons are to become engaged. Since initially learned representations have many more engaged neurons, while representations explored by drift have far fewer engaged neurons, such asymmetrical error histograms are relatively more common for learned representations than for representations explored by drift. Thus if initially learned solution produces an asymmetrical error histogram, drift tends to remove this bias by shifting the mean of the histograms closer to 0 (Suppl. Fig. 5D,E).

### C. Relation to Biology/Experiments

#### C.2. Decorrelation of synaptic weights and neural representations over time

The rate of representational turnover may change over time. As we model drift as diffusion in the synaptic weight solution space, synaptic weights at neighboring time points are correlated. Over longer periods, this correlation gradually decreases and equilibrates (Suppl. Fig. 6A). The decorrelation of synaptic weights corresponds to a parallel decorrelation of neural representations over time (Suppl. Fig. 6B). However, not all synaptic weight changes result in changes to neural activity. Of the three types of flexibility—unconstrained dimensions, semi-constrained dimensions, and drift dimensions—only the drift dimensions driven by engaged neurons lead to representational changes. While learning generates densely engaged representations, early drift induces sparsification, transforming drift dimensions into semi-constrained dimensions. Consequently, early in the drift process, most synaptic changes produce representational changes. In contrast, after equilibration, when many neurons are disengaged, more synaptic changes occur without affecting representations, reducing the rate of representational change. In our simulations, diffusion in the weight space results in comparable rates of synaptic weight changes during both early and later drift (Suppl. Fig. 6C). However, representations change more rapidly during early drift than later drift (Suppl. Fig. 6D). This phenomenon is consistent with experimental findings, as the rate of representational changes shows an initial decrease. While it is unclear whether the greater initial representational changes are solely due to learning or also to drift, our model predicts that even after learning is completed, the rate of representational drift may differ, with greater drift occurring soon after learning and less drift later.

**Supplementary Figure 5.**
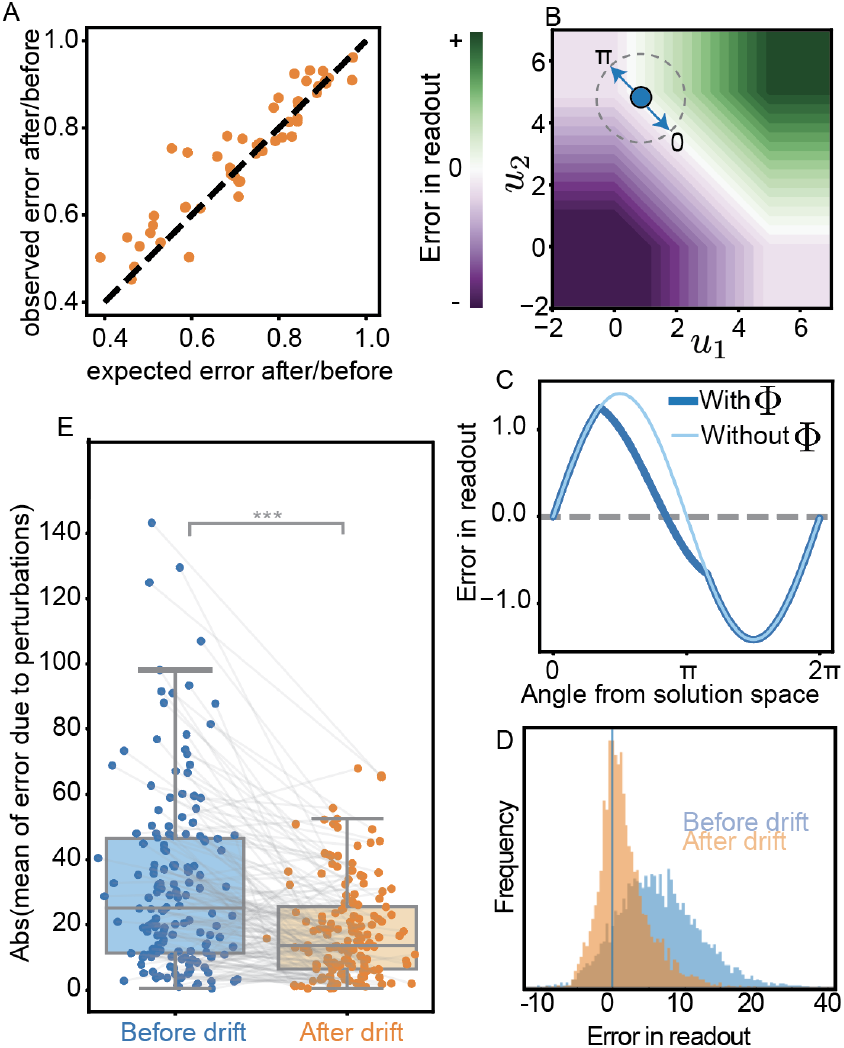
Understanding the robustness of solutions achieved by drift. (A) The x-axis represents 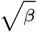, the expected ratio of errors after drift to before drift, while the y-axis shows the observed fraction of errors after drift compared to before drift for each of the 50 simulations from Fig. 4B). (B) Toy example with *z*_*µ*_ = 6. Unsigned error in readout *z* for various (*u*_1_, *u*_2_) values with a schematic demonstrating weight perturbations in two opposite directions. The arrow towards *π* shows perturbation in the direction of threshold crossing. (C) Unsigned error in readout due to a perturbation of size 1 from densely engaged solution (*u*_1_, *u*_2_) = (1.3, 4.7) with the 2nd neuron’s activity *y*_2_ = 4.7 very close to the saturation threshold *α* = 5. The dark blue line indicates unsigned errors in a regular network with Φ non-linearity showing an asymmetry with less positive errors and more negative errors. While the light blue line indicates symmetric errors in the absence of non-linearity. (D) Histogram of errors due to weight perturbations *dU* ∼ *𝒩* (0, 1) in blue for the initially learned solution, and in orange for the solution after drift, showing mean centering of the error histogram through drift. (E) Mean of unsigned error due to weight perturbations for each of the 8 input conditions for 20 different simulations before drift and after drift. n=160, p<0.001***, Wilcoxon test.

#### C.2. Exploring representations with many inactive neurons through drift

Drift can also favor sparsely active representations. Previously, we drew inputs *X* from 𝒩 (0, 1), allowing for both positive and negative values, which resulted in input currents being equally likely to be positive or negative. Additionally, we simulated representational drift in networks with no bias or baseline activity (*B* was a zero matrix in *Z* = *W* Φ (*UX* + *B*)) and set the activity threshold at 0. Under these conditions, the balance of positive and negative input currents resulted in neurons being equally likely to be inactive or active (engaged or saturated). Since we allowed very large weight norms, the exploration range was greater for saturated neurons compared to engaged neurons. Consequently, most active neurons were saturated, and the representations were sparsely engaged, consisting predominantly of inactive and saturated neurons. However, adjusting these parameters can reduce the likelihood of saturated neurons. For instance, when inputs *X* are restricted to non-negative values and paired with a negative bias or baseline, the exploration range for inactive neurons increases, while that for saturated neurons decreases. Additionally, increasing the saturation threshold *α* and imposing smaller weight norm bounds further reduce the prevalence of representations with saturated neurons.

To favor inactive neurons, we configured *B* as a constant matrix with a value of -15, initialized weights from *U* 𝒩(0, 0.5), enforced smaller weight norm bounds, and defined the activity and saturation thresholds as 0 and 10, respectively. This configuration allowed drift to extensively explore large negative input currents (Suppl. Fig. 7A) with smaller changes in weight distribution (Suppl. Fig. 7B). While most neurons were active before drift, only a few remained active afterward (Suppl. Fig. 7C,D). These representational changes occurred despite stable readouts (Suppl. Fig. 7E). Drift led to an increase and eventual plateau in the number of inactive neurons, a decrease in the number of engaged neurons, and a consistently low number of saturated neurons (Suppl. Fig. 7F,G) producing sparsely active representations more akin to biological systems. Like sparsely engaged solutions, these sparsely active representations also showed robustness to synaptic weight changes *U* (Suppl. Fig. 7H). In addition, they also showed robustness to changes in readout weights *W* as error due to *W* changes is proportional to activity levels (Suppl. Fig. 7I). In these simulations, we observed a variety of single-neuron behaviors such as intermittent firing (Suppl. Fig. 7J, input condition 7), relatively stable firing at a particular input condition (Suppl. Fig. 7K, input condition 6), field switching between input conditions (Suppl. Fig. 7K, input conditions 2 and 3), and gain or loss of fields (Suppl. Fig. 7L, input conditions 0 and 5). Neurons also exhibited heterogeneous firing rates: some cells fired at high rates across multiple input conditions (Suppl. Fig. 7K), while others that remained less active (Suppl. Fig. 7J,L).

**Supplementary Figure 6.**
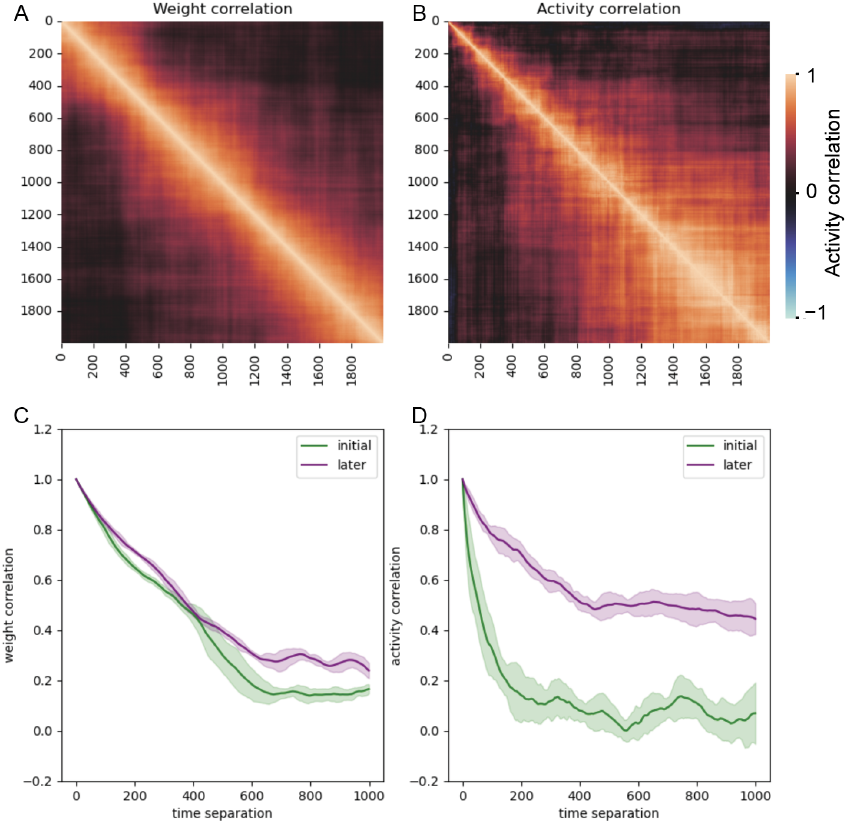
Decorrelation of synaptic weights and neural representation. (A). Heatmap of the correlation between synaptic weights at different timepoints during drift, illustrating a gradual decorrelation over time. (B) Heatmap of the correlation between neural representations for all 8 input conditions at different timepoints, showing an initial rapid decorrelation followed by a relatively slower rate of decorrelation. (C) Lines depict the correlation between average synaptic weights during the first 10,000 timepoints of early drift (green) and during a 10,000-timepoint interval at a later stage of drift, compared to weights at 100,000 timepoints. Shaded area covers one standard deviation away from the mean. (D) Similar to (C), but showing the decorrelation of neural activity with faster decorrelation during early and a slower rate of decorrelation in the later stages.

#### C.3. Drift helps decorrelate neural representations across different input conditions

Solutions explored by drift can provide additional advantages, such as better separability of neural representations. In biology, neural representations for two different input conditions can be highly correlated when memories are encoded close in time, as the same neurons are excitable and thus may get incorporated into both representations. However, if animals need to distinguish between these two different inputs, correlated representations pose a challenge. They are not easily separable, require carefully chosen decision boundaries for classification, and are not robust to noise. In contrast, orthogonal representations for different input conditions enhance their separability. If learned representations for distinct inputs are correlated, orthogonalizing them could be beneficial. Drift can naturally lead to orthogonal representations because, in high-dimensional spaces, there are many more orthogonal vectors than correlated ones. This implies that the representations explored for different input conditions are likely to become decorrelated. To model this phenomenon, we used a network with a constant bias/baseline of − 15, and initialized the wights with a smaller standard deviation *U*∼ 𝒩 (0, 0.5), to keep the weight norms low and avoid saturated neurons. During allocation, instead of randomly choosing *η* for each *µ*, we generated highly correlated *η*_*µ*_, and then performed gradient descent to learn the mappings from inputs to readouts for each *µ* = 1, …., *P*. Though the inputs were decorrelated (Suppl. Fig. 8A), the learned neural representations for different input conditions *µ* were highly correlated (Suppl. Fig. 8B). Upon drifting we found that the system explored sparsely active representations that became decorrelated across different input conditions (Suppl. Fig. 8C-G), while maintaining fixed readouts and relatively stable weight norms (Suppl. Fig. 8H,I). This suggests that mere random exploration of solution space can help the system by finding decorrelated representations that improve separability.

**Supplementary Figure 7.**
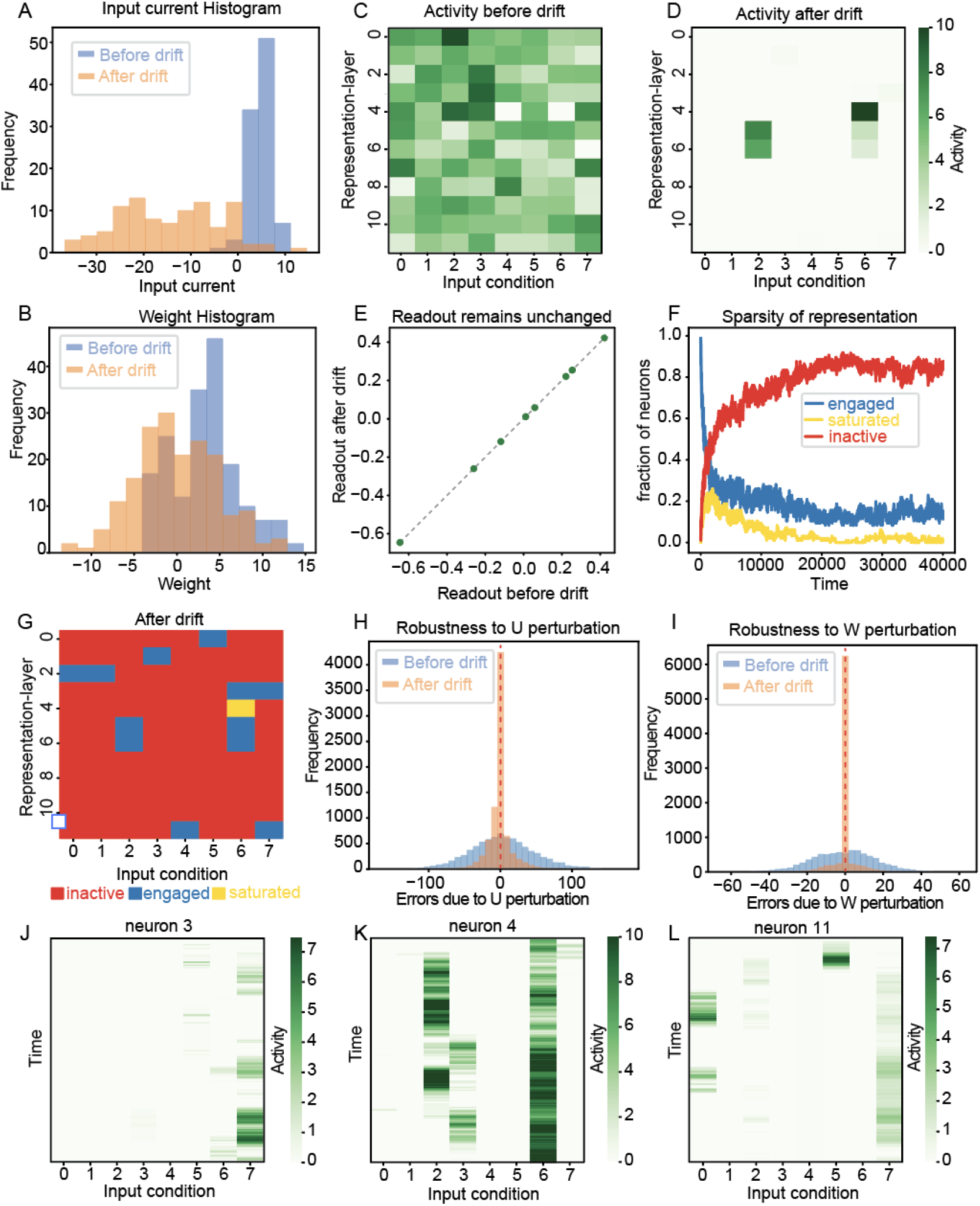
Reaching sparsely active solutions through drift. (A) Histogram of input currents before (blue) and after drift (orange), showing exploration of negative input currents after drift. (B) Histogram of weights before (blue) and after drift (orange) showing stable weight distribution. (C) Neural representation under different input conditions before drift is densely active. (D) Neural representation after drift is sparsely active. (E) Readout before and after drift remains the same. (F) Fraction of engaged, inactive, and saturated neurons during drift, showing increase in the number of inactive neurons. (G) Inactive, engaged, and saturated neurons after drift. (H) Errors due to perturbation of synaptic weights *U*, showing drift improves robustness to U changes. (I) Errors due to perturbation of readout weights *W*, showing drift can also improve robustness to *W* changes when exploring sparsely active solutions. (J,K,L) 3 Example neurons’ activities under different input conditions over time during drift showing relatively stable fields, intermittent firing, gain, loss, and switching of fields.

**Supplementary Figure 8.**
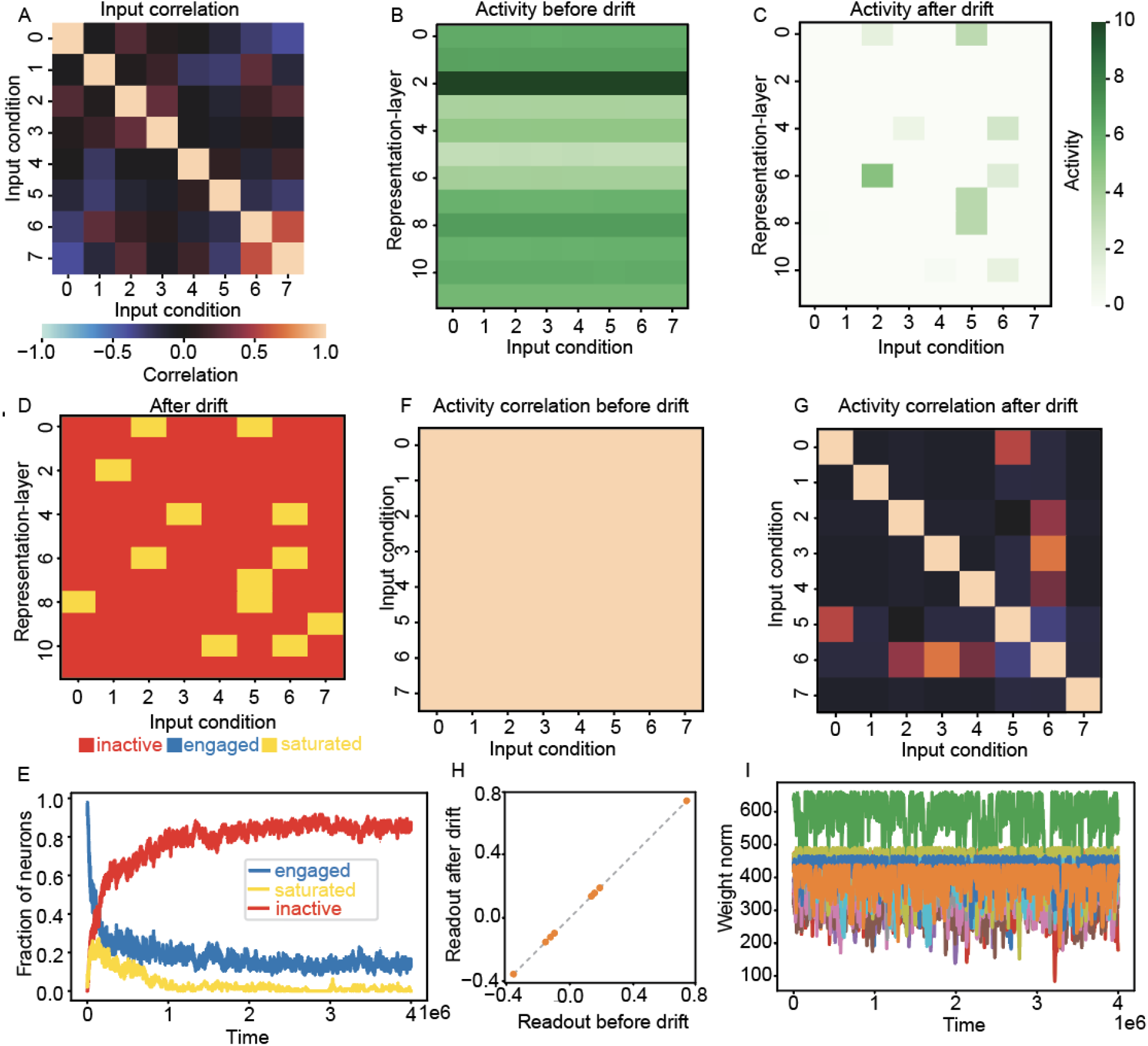
Decorrelation of representations across different input conditions. (A) Correlation across *P* = 8 different input conditions. (B) Neural representations under these *P* = 8 input conditions after gradient descent (before drift), showing they are very similar due to correlated *η* allocation. (C) Sparsely active neural representations after drift. (D) Inactive, engaged, and saturated neurons after drift. (E) Fraction of engaged, inactive, and saturated neurons during drift, showing increase in the number of inactive neurons. (F) Correlation of neural activity across different input conditions after learning, showing the representations are highly correlated. (G) Correlation of neural activity across different input conditions after drift, demonstrating that drift has decorrelated the representations. (H) Readout before and after drift, showing maintenance of readouts. (I) Weight norms for each neuron (different colors) over time during drift, initialized with smaller weights to prevent large input currents.

### D. SI Methods

To curtail synaptic weights *U* from becoming excessively large, we introduced a constraint that limits the magnitude of weights associated with each representation-layer neuron *j*, such that 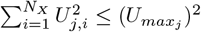, where 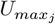 is the maximum weight norm allowed for neuron *j*, which allows heterogeneity across neurons but maintains stability over time. After, every learning event, we set 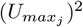to be 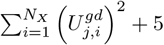, where *U* ^*gd*^ is the learned synaptic weight matrix. When a proposed change *δη* caused the weights onto a representation-layer neuron *j* to exceed its weight norm bound 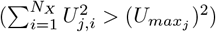, we adjusted the change so that the weight norm exactly equals the bound (*U*_*max*_)^2^.

However, in this process, we still need to ensure that the readouts are preserved. To achieve this, we scaled the proposed change *δη*_*j,µ*_ by *γ*_*j*_ only when *j* ∉ *ℰ* ^*µ*^, so that the activity *Y*_*j,µ*_ and thus the readout *Z*_*µ*_ remain the same. The new *η* at time *t* + 1 is given by

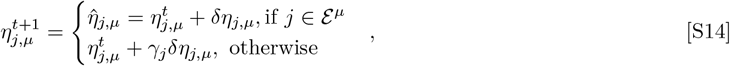

where 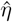 is the new *η* with the proposed change *δη*. Since, changes can only be made in specific *η*_*j,µ*_, we first write the weight norm in terms of 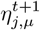

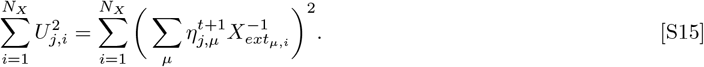

For the updated weights at time *t* + 1, we want the weight norm to be 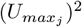Additionally, to target changes specifically in the non-engaged dimensions of *η*, we separate the contribution of *µ*s based on whether *j* ∈ ℰ^*µ*^. As a result we obtain

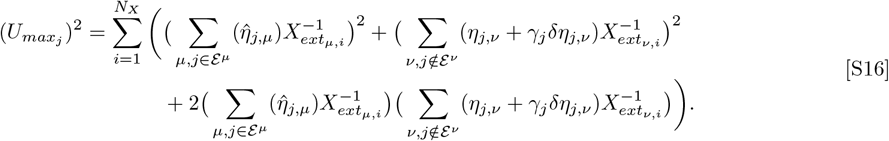

Rearranging terms from S16, we get a quadratic equation in *γ*_*j*_:

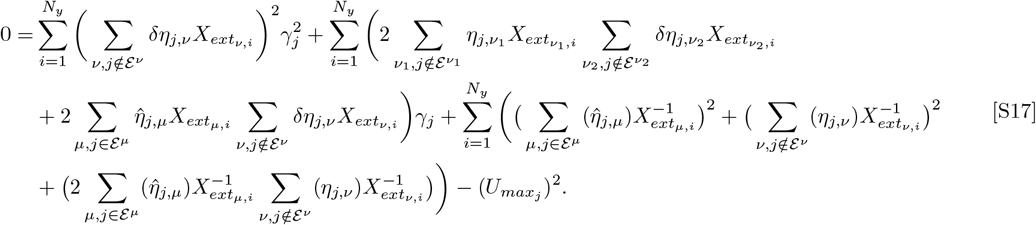

We solve for *γ* using S17, choose a root that satisfies 0 ≤ *γ* ≤ 1, and then find *η*^*t*+1^ using S14.

## Notes

### Competing Interest Statement

The authors have declared no competing interest.

## References

1. DO Hebb, The organization of behavior. (John Wiley and Sons Inc.), (1964).

2. JJ Langille, RE Brown, The synaptic theory of memory: A historical survey and reconciliation of recent opposition. Front. Syst. Neurosci. 12 (2018).

3. W Sun, M Advani, N Spruston, A Saxe, JE Fitzgerald, Organizing memories for generalization in complementary learning systems. Nat. Neurosci. 26, 1438–1448 (2023).

4. CG Kentros, NT Agnihotri, S Streater, RD Hawkins, ER Kandel, Increased attention to spatial context increases both place field stability and spatial memory. Neuron 42, 283–295 (2004).

5. EA Mankin, et al., Neuronal code for extended time in the hippocampus. Proc. Natl. Acad. Sci. 109, 19462–19467 (2012).

6. Y Ziv, et al., Long-term dynamics of ca1 hippocampal place codes. Nat. Neurosci. 16, 264–266 (2013).

7. A Rubin, N Geva, L Sheintuch, Y Ziv, Hippocampal ensemble dynamics timestamp events in long-term memory. eLife 4 (2015).

8. JS Lee, JJ Briguglio, JD Cohen, S Romani, AK Lee, The statistical structure of the hippocampal code for space as a function of time, context, and value. Cell 183 (2020).

9. SJ Levy, NR Kinsky, W Mau, DW Sullivan, ME Hasselmo, Hippocampal spatial memory representations in mice are heterogeneously stable. Hippocampus 31, 244–260 (2020).

10. W Hockeimer, RY Lai, M Natrajan, W Snider, JJ Knierim. Leveraging place field repetition to understand positional versus nonpositional inputs to hippocampal field CA1 (2022).

11. ME Rule, T O’Leary, CD Harvey, Causes and consequences of representational drift. Curr. Opin. Neurobiol. 58, 141–147 (2019).

12. LN Driscoll, L Duncker, CD Harvey, Representational drift: Emerging theories for continual learning and experimental future directions. Curr. Opin. Neurobiol. 76, 102609 (2022).

13. U Rokni, AG Richardson, E Bizzi, HS Seung, Motor learning with unstable neuralnbsp;representations. Neuron 54, 653–666 (2007).

14. AR Chambers, S Rumpel, A stable brain from unstable components: Emerging concepts and implications for neural computation. Neuroscience 357, 172–184 (2017).

15. D Kappel, R Legenstein, S Habenschuss, M Hsieh, W Maass, A dynamic connectome supports the emergence of stable computational function of neural circuits through reward-based learning. eneuro 5 (2018).

16. P Masset, S Qin, JA Zavatone-Veth, Drifting neuronal representations: Bug or feature? Biol. Cybern. 116, 253–266 (2022).

17. S Qin, et al., Coordinated drift of receptive fields in hebbian/anti-hebbian network models during noisy representation learning. Nat. Neurosci. 26, 339–349 (2023).

18. LN Driscoll, NL Pettit, M Minderer, SN Chettih, CD Harvey, Dynamic reorganization of neuronal activity patterns in parietal cortex. Cell 170 (2017).

19. LA DeNardo, et al., Temporal evolution of cortical ensembles promoting remote memory retrieval. Nat. Neurosci. 22, 460–469 (2019).

20. D Deitch, A Rubin, Y Ziv, Representational drift in the mouse visual cortex. Curr. Biol. 31 (2021).

21. HC Wang, AM LeMessurier, D. Feldman, Tuning instability of non-columnar neurons in the salt-and-pepper whisker map in somatosensory cortex. Nat. Commun. 13 (2022).

22. CE Schoonover, SN Ohashi, R Axel, AJ Fink, Representational drift in primary olfactory cortex. Nature 594, 541–546 (2021).

23. A Rubin, et al., Revealing neural correlates of behavior without behavioral measurements. Nat. Commun. 10 (2019).

24. J Xia, TD Marks, MJ Goard, R Wessel, Stable representation of a naturalistic movie emerges from episodic activity with gain variability. Nat. Commun. 12 (2021).

25. AT Keinath, CA Mosser, MP Brandon, The representation of context in mouse hippocampus is preserved despite neural drift. Nat. Commun. 13 (2022).

26. K Aitken, M Garrett, S Olsen, S Mihalas, The geometry of representational drift in natural and artificial neural networks. PLOS Comput. Biol. 18 (2022).

27. DG Wyrick, et al., Differential encoding of temporal context and expectation under representational drift across hierarchically connected areas. bioRxiv (2023).

28. ZN Roth, EP Merriam, Representations in human primary visual cortex drift over time. Nat. Commun. 14 (2023).

29. JL McClelland, BL McNaughton, R. O’Reilly, Why there are complementary learning systems in the hippocampus and neocortex: insights from the successes and failures of connectionist models of learning and memory. Psychol. review 102, 419 (1995).

30. MK Benna, S Fusi, Computational principles of synaptic memory consolidation. Nat. neuroscience 19, 1697–1706 (2016).

31. D Kappel, S Habenschuss, R Legenstein, W Maass, Network plasticity as bayesian inference. PLOS Comput. Biol. 11 (2015).

32. T Biswas, JE Fitzgerald, Geometric framework to predict structure from function in neural networks. Phys. Rev. Res. 4 (2022).

33. T Biswas, TL Li, JE Fitzgerald, Tensor formalism for predicting synaptic connections with ensemble modeling or optimization. arXiv preprint arXiv:2310.20309 (2023).

34. JJ DiCarlo, DD Cox, Untangling invariant object recognition. Trends cognitive sciences 11, 333–341 (2007).

35. T Biswas, WE Bishop, JE Fitzgerald, Theoretical principles for illuminating sensorimotor processing with brain-wide neuronal recordings. Curr. opinion neurobiology 65, 138–145 (2020).

36. ME Rule, et al., Stable task information from an unstable neural population. eLife 9 (2020).

37. E Chan, O Baumann, MA Bellgrove, JB Mattingley, From objects to landmarks: The function of visual location information in spatial navigation. Front. Psychol. 3 (2012).

38. RM Seyfarth, DL Cheney, P Marler, Monkey responses to three different alarm calls: Evidence of predator classification and semantic communication. Science 210, 801–803 (1980).

39. W Mau, ME Hasselmo, DJ Cai, The brain in motion: How ensemble fluidity drives memory-updating and flexibility. eLife 9 (2020).

40. Y Tian, Y Zhang, H Zhang, Recent advances in stochastic gradient descent in deep learning. Mathematics 11, 682 (2023).

41. A Ratzon, D Derdikman, O Barak, Representational drift as a result of implicit regularization. bioRxiv (2023).

42. S Káli, P Dayan, Off-line replay maintains declarative memories in a model of hippocampal-neocortical interactions. Nat. Neurosci. 7, 286–294 (2004).

43. S Grossberg, How does a brain build a cognitive code? Psychol. Rev. 87, 1–51 (1980).

44. M Mermillod, A Bugaiska, P Bonin, The stability-plasticity dilemma: Investigating the continuum from catastrophic forgetting to age-limited learning effects. Front. Psychol. 4 (2013).

45. A Attardo, JE Fitzgerald, MJ Schnitzer, Impermanence of dendritic spines in live adult ca1 hippocampus. Nature 523, 592–596 (2015).

46. NE Ziv, N Brenner, Synaptic tenacity or lack thereof: spontaneous remodeling of synapses. Trends neurosciences 41, 89–99 (2018).

47. YF Kalle Kossio, S Goedeke, C Klos, RM Memmesheimer, Drifting assemblies for persistent memory: Neuron transitions and unsupervised compensation. Proc. Natl. Acad. Sci. 118 (2021).

48. F Pashakhanloo, A Koulakov, Stochastic gradient descent-induced drift of representation in a two-layer neural network. arXiv (2023).

49. G Delamare, Y Zaki, DJ Cai, C Clopath, Drift of neural ensembles driven by slow fluctuations of intrinsic excitability. eLife (2023).

50. W Sun, et al., Learning produces a hippocampal cognitive map in the form of an orthogonalized state machine. bioRxiv pp. 2023–08 (2023).

51. T Faber, J Joerges, R Menzel, Associative learning modifies neural representations of odors in the insect brain. Nat. Neurosci. 2, 74–78 (1999).

52. SW Failor, M Carandini, KD Harris. Visuomotor Assoc. orthogonalizes visual cortical population codes (2021).

53. D George, et al., Clone-structured graph representations enable flexible learning and vicarious evaluation of cognitive maps. Nat. communications 12, 2392 (2021).

54. NA Cayco-Gajic, RA Silver, Re-evaluating circuit mechanisms underlying pattern separation. Neuron 101, 584–602 (2019).

55. R Ajemian, A D’Ausilio, H Moorman, E Bizzi, A theory for how sensorimotor skills are learned and retained in noisy and nonstationary neural circuits. Proc. Natl. Acad. Sci. 110 (2013).

56. S Ganguli, H Sompolinsky, Compressed sensing, sparsity, and dimensionality in neuronal information processing and data analysis. Annu. review neuroscience 35, 485–508 (2012).

57. JB Aimone, J Wiles, FH Gage, Computational influence of adult neurogenesis on memory encoding. Neuron 61, 187–202 (2009).

58. DJ Cai, et al., A shared neural ensemble links distinct contextual memories encoded close in time. Nature 534, 115–118 (2016).

59. A Kafkas, D Montaldi, How do memory systems detect and respond to novelty? Neurosci. letters 680, 60–68 (2018).

60. M Zhu, SJ Kuhlman, A. Barth, Transient enhancement of stimulus-evoked activity in neocortex during sensory learning. Learn. amp; Mem. 31 (2024).

61. Q Lu, U Hasson, KA Norman, A neural network model of when to retrieve and encode episodic memories. elife 11, e74445 (2022).

62. RS Blumenfeld, C Ranganath, Prefrontal cortex and long-term memory encoding: an integrative review of findings from neuropsychology and neuroimaging. The Neurosci. 13, 280–291 (2007).

63. GG Turrigiano, The self-tuning neuron: synaptic scaling of excitatory synapses. Cell 135, 422–435 (2008).

64. Q Hou, D Zhang, L Jarzylo, RL Huganir, HY Man, Homeostatic regulation of ampa receptor expression at single hippocampal synapses. Proc. Natl. Acad. Sci. 105, 775–780 (2008).

65. JC Béïque, Y Na, D Kuhl, PF Worley, RL Huganir, Arc-dependent synapse-specific homeostatic plasticity. Proc. Natl. Acad. Sci. 108, 816–821 (2011).

66. N Vitureira, M Letellier, Y Goda, Homeostatic synaptic plasticity: from single synapses to neural circuits. Curr. opinion neurobiology 22, 516–521 (2012).

67. SA Josselyn, PW Frankland, Memory allocation: mechanisms and function. Annu. review neuroscience 41, 389–413 (2018).

68. AJ Silva, Y Zhou, T Rogerson, J Shobe, J Balaji, Molecular and cellular approaches to memory allocation in neural circuits. Science 326, 391–395 (2009).

69. G Dragoi, S Tonegawa, Preplay of future place cell sequences by hippocampal cellular assemblies. Nature 469, 397–401 (2011).

70. GI Parisi, R Kemker, JL Part, C Kanan, S Wermter, Continual lifelong learning with neural networks: A review. Neural networks 113, 54–71 (2019).

71. AL Barth, JF Poulet, Experimental evidence for sparse firing in the neocortex. Trends Neurosci. 35, 345–355 (2012).

72. C Chen, et al. Single field evolution rule governs dynamics representational drift mouse hippocampal dorsal ca1 region (2024).

73. D Attwell, SB Laughlin, An energy budget for signaling in the grey matter of the brain. J. Cereb. Blood Flow amp; Metab. 21, 1133–1145 (2001).

74. S Laughlin, Energy as a constraint on the coding and processing of sensory information. Curr. Opin. Neurobiol. 11, 475–480 (2001).

75. G Wang, R Wang, W Kong, J Zhang, The relationship between sparseness and energy consumption of neural networks. Neural Plast. 2020, 1–13 (2020).

76. HH Kornhuber, Neural control of input into long term memory: Limbic system and amnestic syndrome in man. Mem. Transf. Inf. p. 1–22 (1973).

77. G Delamare, DF Tomé, C Clopath, Intrinsic neural excitability biases allocation and overlap of memory engrams. The J. Neurosci. 44 (2024).

78. T Rogerson, et al., Synaptic tagging during memory allocation. Nat. Rev. Neurosci. 15, 157–169 (2014).

79. D Nitz, B McNaughton, Differential modulation of ca1 and dentate gyrus interneurons during exploration of novel environments. J. Neurophysiol. 91, 863–872 (2004).

80. J Csicsvari, J O’Neill, K Allen, T Senior, Place-selective firing contributes to the reverse-order reactivation of ca1 pyramidal cells during sharp waves in open-field exploration. Eur. J. Neurosci. 26, 704–716 (2007).

81. D Khatib, et al., Active experience, not time, determines within-day representational drift in dorsal ca1. Neuron (2023).

82. MP Karlsson, LM Frank, Network dynamics underlying the formation of sparse, informative representations in the hippocampus. The J. Neurosci. 28, 14271–14281 (2008).

83. L Sheintuch, N Geva, D Deitch, A Rubin, Y Ziv, Organization of hippocampal ca3 into correlated cell assemblies supports a stable spatial code. Cell Reports 42, 112119 (2023).

84. TD Marks, MJ Goard, Stimulus-dependent representational drift in primary visual cortex. Nat. Commun. 12 (2021).

85. S Qin, C Pehlevan, Representational sparsity determines representational stability in sensory cortices. (year?).

86. G Elyasaf, A Rubin, Y Ziv, Novel off-context experience constrains hippocampal representational drift. Curr. Biol. (2024).

87. SP Vaidya, G Li, RA Chitwood, Y Li, JC Magee. The formation an expanding memory representation hippocampus (2023).

88. C van Vreeswijk, H Sompolinsky, Chaos in neuronal networks with balanced excitatory and inhibitory activity. Science 274, 1724–1726 (1996).

89. B Haider, A Duque, AR Hasenstaub, D. McCormick, Neocortical network activityin vivois generated through a dynamic balance of excitation and inhibition. The J. Neurosci. 26, 4535–4545 (2006).

90. M Valero, A Navas-Olive, LM de la Prida, G Buzsáki, Inhibitory conductance controls place field dynamics in the hippocampus. Cell Reports 40, 111232 (2022).

91. S Crochet, J Poulet, Y Kremer, C Petersen, Synaptic mechanisms underlying sparse coding of active touch. Neuron 70, 170 (2011).

92. M Rudolph, M Pospischil, I Timofeev, A Destexhe, Inhibition determines membrane potential dynamics and controls action potential generation in awake and sleeping cat cortex. The J. Neurosci. 27, 5280–5290 (2007).

93. O Amsalem, H Inagaki, J Yu, K Svoboda, R Darshan, Sub-threshold neuronal activity and the dynamical regime of cerebral cortex. Nat. Commun. 15 (2024).

94. JH Kim, K Daie, N Li, A combinatorial neural code for long-term motor memory. Nature (2024).

